# Serotonin predictively encodes value

**DOI:** 10.1101/2023.09.19.558526

**Authors:** Emerson F. Harkin, Cooper D. Grossman, Jeremiah Y. Cohen, Jean-Claude Béïque, Richard Naud

**Affiliations:** Department of Cellular and Molecular Medicine, University of Ottawa, K1H 8M5, Ottawa, Canada; Centre for Neural Dynamics, University of Ottawa, K1H 8M5, Ottawa, Canada; Brain and Mind Institute, University of Ottawa, K1H 8M5, Ottawa, Canada; California Institute of Technology, Pasadena, United States; Allen Institute for Neural Dynamics, 98109, Seattle, United States; Department of Physics, University of Ottawa, K1H 8M5, Ottawa, Canada

**Keywords:** serotonin, reinforcement learning, neural coding

## Abstract

The *in vivo* responses of dorsal raphe nucleus (DRN) serotonin neurons to emotionally-salient stimuli are a puzzle. Existing theories centred on reward, surprise, or uncertainty individually account for some aspects of serotonergic activity but not others. Here we find a unifying perspective in a biologically-constrained predictive code for cumulative future reward, a quantity called state value in reinforcement learning. Through simulations of trace conditioning experiments common in the serotonin literature, we show that our theory, called value prediction, intuitively explains phasic activation by both rewards and punishments, preference for surprising rewards but absence of a corresponding preference for punishments, and contextual modulation of tonic firing—observations that currently form the basis of many and varied serotonergic theories. Next, we re-analyzed data from a recent experiment and found serotonin neurons with activity patterns that are a surprisingly close match: our theory predicts the marginal effect of reward history on population activity with a precision ≪0.1 Hz neuron^−1^. Finally, we directly compared against quantitative formulations of existing ideas and found that our theory best explains both within-trial activity dynamics and trial-to-trial modulations, offering performance usually several times better than the closest alternative. Overall, our results show that previous models are not wrong, but incomplete, and that reward, surprise, salience, and uncertainty are simply different faces of a predictively-encoded value signal. By unifying previous theories, our work represents an important step towards understanding the potentially heterogeneous computational roles of serotonin in learning, behaviour, and beyond.

## Introduction

What do the activity patterns of serotonin neurons encode? Over a quarter-century ago, Schultz, Dayan, and Montague (1) persuasively argued that the phasic activity of dopamine neurons might encode the reward prediction errors (RPEs) of reinforcement learning (RL) theory (2). Given the deep connections between the dopamine and serotonin systems, both of which are neuromodulatory systems with important and well-studied roles in regulating mood, learning, and behaviour (3), it is surprising that no single account of the responses of serotonin neurons enjoys a similar level of support.

There are several possible reasons for this lack of consensus. One possibility is that the serotonin system is not a monolith, but rather a heterogeneous collection of partially-overlapping sub-systems with diverse coding features (4–6). Another possibility, in no way mutually-exclusive, arises from the fact that experimental and theoretical work in the serotonin field, including our own, has been deeply shaped by the potentially incorrect assumption that the activity patterns of serotonin neurons can be divided into phasic and tonic components that reflect essentially unrelated quantities (3, 7–10). This separation of timescales is reflected in the currently fragmented picture of the dominant tuning features of serotonin neurons. Rejecting this assumption could lead to more clarity about serotonergic function within and across raphe sub-systems.

Over the past decade, detailed experimental characterizations of the diverse *in vivo* responses of genetically-identified serotonin neurons to emotionally-salient stimuli have revealed some common themes in their tuning features, even if these patterns remain difficult to interpret. In trace conditioning experiments, the activity patterns of serotonin neurons are dominated by phasic bumps, plateaus, or ramps preceding expected rewards that emerge over the course of learning and diminish during reversals (Fig. 1A; 5, 11–13), modulation of tonic activity by reward or punishment context (Fig. 1B; 11), a phasic preference for unpredicted over predicted rewards (Fig. 1C; 11, 13), and phasic activation by punishments whether predicted or not (Fig. 1D; 5, 11, 13). To explain aspects of these observations, serotonin neurons have been proposed to encode current or future reward [Fig. 1A (14) or, separately, B (11), but not D], surprise [Fig. 1C but not D (13)], or salience [Figure 1D but not C (5)]. The reward (14), surprise (13), uncertainty (15), and salience (5) theories do not offer detailed predictions about the dynamics of serotonin neuron activity, nor can any of them individually account for all of their dominant tuning features (Table 1). Other serotonergic theories related to persistence (16), confidence (17), learning rate (18), and discounting (19, 20) focus on explaining the effects of serotonergic manipulations on behaviour and do not connect directly to the naturalistic tuning features of these cells (but see 15). Even the best established tuning features of serotonin neurons therefore lack a consistent interpretation.

**Table 1.** Short timescale reward tuning features of genetically-identified mouse DRN serotonin neurons qualitatively explained by various theories. ✓ and ✗ indicate empirical observations that are clearly consistent or inconsistent with each theory; ambiguous cases are left blank. Ambiguity is due to a lack of quantitative models to accompany the current and future reward, salience, and surprise theories, as well as variation in experimental design to a lesser extent. Note that refs. (12, 14, 38, 39) focus on a signal that qualitatively reflects both current and future reward. This signal is referred to as reward or beneficialness by the authors, but is most similar to a value signal in our terminology.

| Theory | Tonic firing tracks rew. rate | Rew. cue activation (11–13, 15, 38, 39) | Rew. delivery activation (4, 5, 11–13, 38, 39) | Pun. activation (4, 5, 11, 13) | Correlated rew. cue and pun. activation (11) | Surprise rew. preference (11, 13) | Lack of surprise pun. preference (11, 13) |
| --- | --- | --- | --- | --- | --- | --- | --- |
| Value prediction (ours) | ✓(Fig. 3C) | ✓(Fig. 3A) | ✓(Fig. 3A) | ✓(Fig. 3B) | ✓(Fig. 4C) | ✓(Fig. 4B) | ✓(Fig. 4B) |
| Current and future reward (12, 14, 38, 39) | ✓ | ✓ | ✓ | ✗ | ✗ | ✗ | ✗ |
| Salience (5) | ✗ | ✓ | ✓ | ✓ |  | ✗ | ✓ |
| Surprise (13) | ✗ | ✓ | ✓ | ✓ |  | ✓ | ✗ |
| Dopamine opponent (7) | ✓ | ✗ | ✗ | ✓ | ✗ | ✗ | ✗ |

**Fig. 1.**
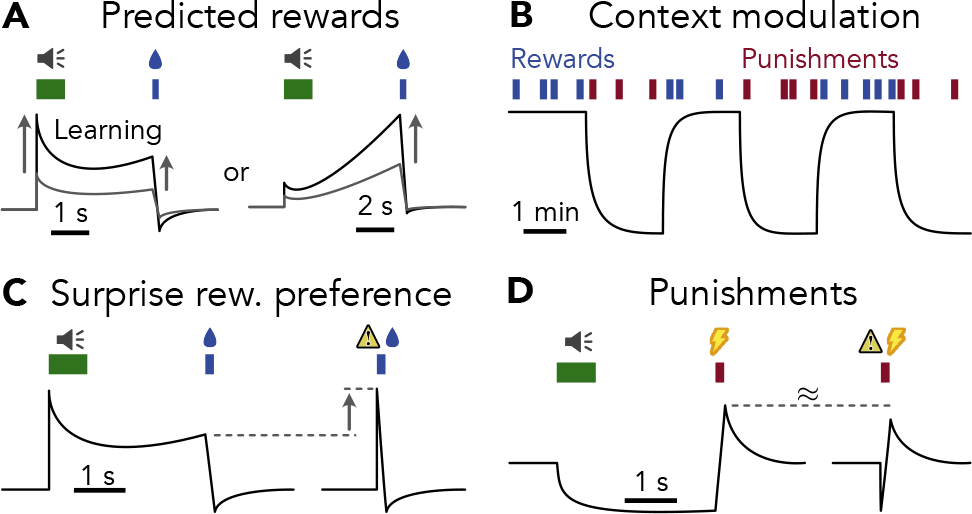
Summary of qualitative tuning features of serotonin neurons captured by predictive value coding model. Curves indicate the activity of serotonin neurons over time, measured either as firing rate (11) or calcium fluorescence (5, 12, 13). **A** Phasic activation by predicted rewards over short timescales (11) emerges gradually during learning (5, 12, 13). Depending on the experiment, activity takes the form of a phasic cue-associated peak followed by a plateau (left; 11, 13), or a ramp leading up to reward (right; (12)). **B** Tonic activity modulated by reward or punishment context over long timescales (11). **C** Stronger phasic activation by unpredicted (right) compared with predicted (left) rewards (11, 13). **D** Phasic activation by punishments whether predicted (left) or not (right) (5, 11, 13).

Here we argue that existing qualitative serotonergic theories are incomplete, not incorrect, and that reward, surprise, salience, and uncertainty are simply different aspects of a single quantity encoded in the activity patterns of serotonin neurons. To formulate a consistent interpretation of the dominant reward and punishment tuning features of serotonin neurons outlined above, we combine top-down ideas from theories of RL (2) and predictive coding (21, 22) with recent bottom-up insights into the computational features of the DRN (10). We hypothesized that serotonin neurons predictively encode a weighted average of future reward, a quantity referred to as state value in RL, via the dominant biophysical feature of this cell type: exceptionally strong and long-lasting spike frequency adaptation. We formalize this hypothesis in a quantitative model that we refer to as as the value prediction theory of serotonin.

To test our value prediction theory, we simulate trace conditioning experiments common in the serotonin literature (11–13, 15) and show that our model provides a consistent account of the main established tuning features of these cells. We also interpolate and extend previous results, providing intuitive mechanistic connections between seemingly unrelated observations, resolving apparent conflicts in the literature, and making experimentally testable predictions. Next, we re-analyze a recently-published dataset from a trace conditioning experiment (23), finding activity patterns consistent with our hypothesis. Finally, to counter our own confirmation bias, we explicitly compare against quantitative formulations of previous theories and find that value prediction best explains the data by a large margin. Our theoretical and empirical results reveal a surprisingly precise quantitative code for value in the serotonin system.

## Results

### Predictive encoding of value signals

Reinforcement learning describes the process by which an agent learns a policy for controlling the state of its environment *S, s* in order to maximize reward *R, r* (Figure 2A; 2). For example, a mouse learning which lever to press to obtain a food pellet in an operant conditioning experiment. RL conceptualizes the reward estimate as a mapping from states to future rewards referred to as a value function *v*(*s*) (Figure 2B). For simplicity, here we focus on the state value *v*(*s*) which is equivalent to the average *q*(*s, a*) value of actions *a* available in state *s* (Appendix A).

**Fig. 2.**
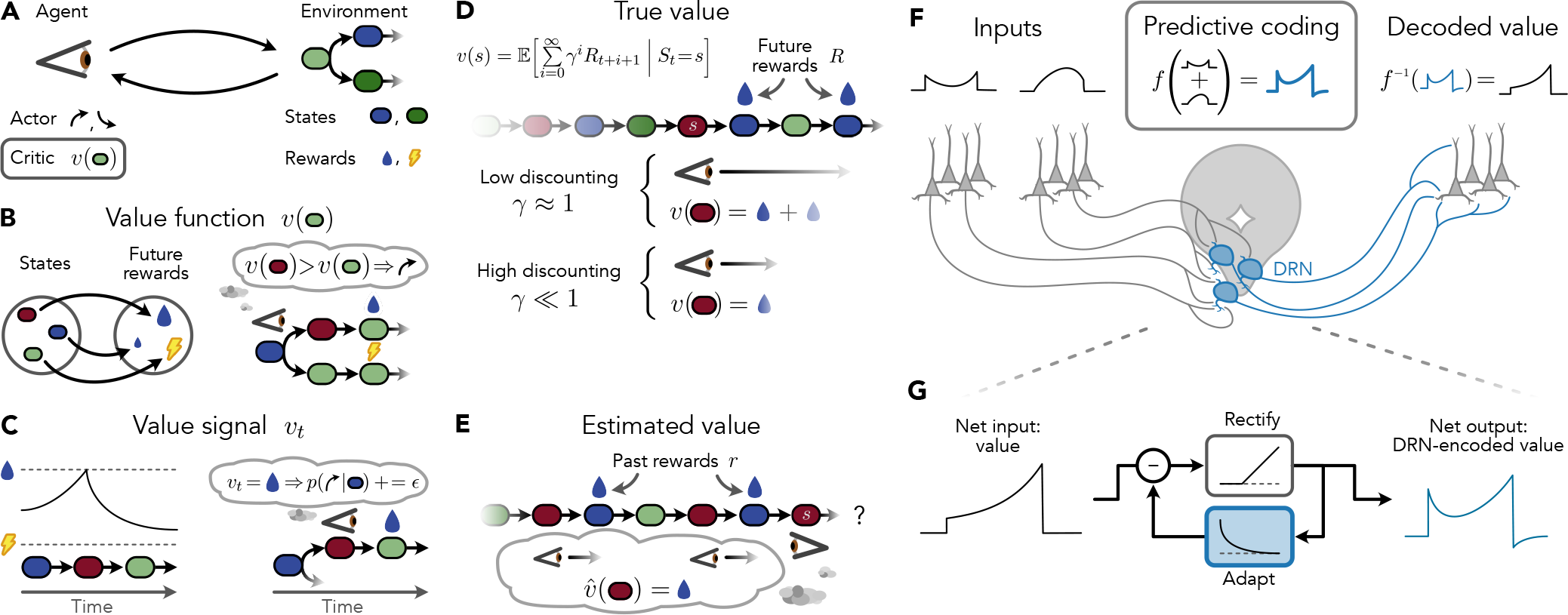
Computing future reward. **A** High-level overview of reinforcement learning (RL; 2). **B** A value function *v*(*s*) is a mapping from states to future rewards. The value function can be used to drive decision-making (right): by comparing the values of future states, the agent can make choices that lead to rewards (24, 25). **C** A value signal *v*_*t*_ is the result of evaluating the value function *v*(*s*) at the current state *s*_*t*_ over time. The value signal can be used to drive learning (right): by increasing the probability of taking an action in proportion to the value signal just after the action is taken, the agent can learn to take actions that lead to rewards (26). **D** Normative definition of a value function as the expected sum of discounted future rewards, referred to as the true value *v*(*s*). **E** The estimated value function 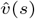 approximates the true value on the basis of past rewards. **F** Predictive value coding model. The dorsal raphe nucleus (DRN) receives a distributed value signal as input, summates its components, and predictively-encodes the result. (Note that although a distributed value code is illustrated, similar to Fig. 2 in ref. 1, it is also possible that the value signal originates in a single upstream region.) The predictively-encoded signal consists of a mixture of the original value signal and its time derivative 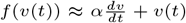, a transformation implemented by strong spike-frequency adaptation in serotonin neurons (10). Predictive coding can easily be reversed via leaky integration in downstream regions to recover the original value signal (Appendix H). **G** Adaptation-based predictive coding model. See Methods.

Value functions have been central to RL since the very beginning (24). One particularly well-known use of value functions is to compare the estimated rewards associated with different hypothetical courses of action (for example, pressing different levers) in order to select the action most likely to lead to the greatest reward (Figure 2B right; 27, 28). A lesser-known application is the evaluation of the current state *s*_*t*_ over time, yielding what we call a value *signal v*_*t*_ = *v*(*s*_*t*_) (Figure 2C). Such a value signal is time dependent because the state is continually changing, a feature exploited by temporal difference learning to gradually refine the estimated values of past states (29, 30). Apart from value learning, value signals can be used to directly reinforce recent actions (26, 31) or promote persistence (27, 32). Here we present evidence of a close match between the activity patterns of serotonin neurons and value, leaving the question of how value might be used to drive learning and behaviour for future work.

Specifically, we focus on a value signal defined as the expected total discounted future reward in the present state

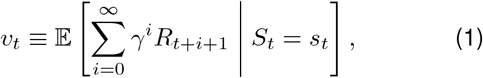

where *R*_*t*_ is the random reward obtained at time *t*, 0 *≤ γ ≤* 1 is a discrete time discounting factor that controls the relative weighting of imminent and distant rewards (imminent rewards are weighted more heavily when *γ* is closer to zero), and 𝔼 [*X*|*Y* = *y*] denotes conditional expectation. Intuitively, it represents a weighted average of future rewards, with closer rewards being weighted more heavily depending on the degree of discounting (Figure 2D). The size of the window within which future rewards are summed to calculate the value can be quantified with the discounting timescale *τ*, defined as *τ* = *−dt/* ln *γ*, where *dt* is the duration of a discrete time step.

The above definition of a value signal in terms of future reward is precise, extremely general (Appendix A), and conceptually simple, but unrealistic: animals do not generally have perfect knowledge of future rewards. We therefore distinguish between signals calculated on the basis of future rewards, which we refer to as *true* value (Figure 2D), and more realistic ones learned from past experience, which we refer to as *estimated* value (Figure 2E).

We propose that the firing rates of serotonin neurons present a predictive code for an estimated value signal (Figure 2F). Recently, we showed that potent spike-frequency adapation dominates the signal processing features of the DRN (10). This removes the part of the signal that is similar to past output, a redundancy-reduction scheme sometimes called predictive coding (21, 22, see 33 for review). In a simplification of this previous work, here we model the firing rate output of the DRN as

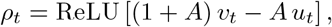

where ReLU[*x*] is the rectified linear function used to ensure the firing rate is non-negative, *A* is a parameter controling the strength of adaptation, and *u*_*t*_ is the adaptation variable, which has a subtractive effect on the output firing rate. We model adaptation as an exponential moving average of past activity (Figure 2G and methods). To build an intuition for this model, consider that, from a computational perspective, adaptation can be seen as implementing temporal differentiation (10, 34). As a consequence, the firing rates of serotonin neurons reflect a mixture of value and its rate of change.

Our main result is that this predictive coding process induces a qualitative change in the value signal. Adaptation has the effect of exaggerating sharp transients, often leading the encoded signal to over- or under-shoot its apparent target (illustrated schematically in Figure 2G, simulations in the next section), and hiding the connection between serotonergic activity patterns and value.

### Value prediction during trace conditioning

To examine the temporal evolution of this signal in an experimental setup common in the serotonin literature, we simulated our model under trace conditioning. Trace conditioning experiments consist of a series of trials that begin with a sensory cue (e.g., an auditory tone or an odour) and end in a reward (typically a drop of water) with a short delay (*∼*2 s) separating the two (11–13). In this experimental paradigm, true value signals take four distinct phases (see Methods): 1) jumping to a higher value upon receiving the cue since the cue indicates a reward is coming, 2) ramping upward between the time of the sensory cue and the reward delivery due to the effects of time discounting, 3) falling during the reward epoch as the *future* reward left to collect disappears, and 4) staying at a constant non-zero value during the inter-trial interval as the animal waits for the next randomly-timed trial to begin (black line in Figure 3A).

**Fig. 3.**
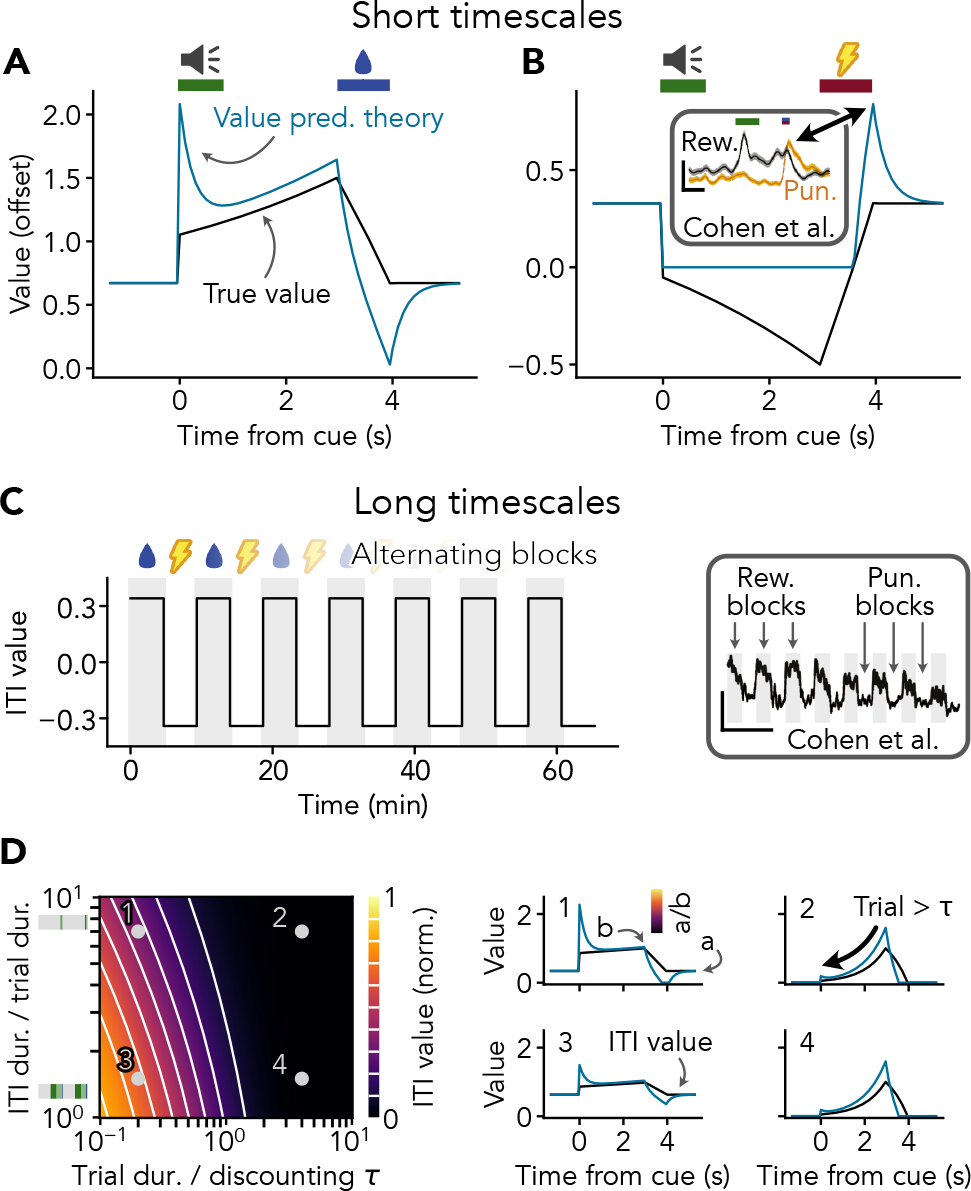
Value prediction signals reward and punishment over multiple timescales. **A**,**B** True value signals (black) and their predictively-encoded counterparts (blue) for trace conditioning trials terminating in either a reward (A) or punishment (B). Signals are shifted up by 0.5 AU to capture background firing. Note resemblance between value prediction theory and firing rate of a genetically-identified serotonin neuron from Cohen *et al*. (11) (B inset; modified from ref. Fig. 3A2; scale bar 5 Hz, 1 s). **C** ITI value reflects reward or punishment context. Simulated block length of 10 trials, normalized reward and punishment sizes of 1 and *−*1, respectively, and all other parameters as in D1 below. Note resemblance with the tonic firing rate of a serotonin neuron (right; modified from ref. 11 Fig. 3B1; scale bar 3 Hz, 10 min). **D** Analytically-derived true value of the inter-trial interval (ITI) is proportional to peak within-trial value. Heatmap shows ITI value as a function of experimental design parameters (mean ITI duration / trial duration; vertical axis; ribbons are to scale, gray represents ITI duration and colours represent trial epochs) and agent parameters (trial duration / discounting *τ* ; horizontal axis). ITI value is presented as a fraction of the peak value during the trial (*i*.*e*., value just before reward). Numbered panels at right illustrate the within-trial dynamics of the true (black) and predictively-encoded (blue) value signals for various combinations of experimental and agent parameters indicated on the heatmap. Note that predictive coding has no effect on ITI value because the time derivative of the value signal during the ITI is zero. Since value signals are normalized, different reward sizes can be accommodated by scaling traces. Trial structure same as in A. See Figure 8 for an extended range of trial durations and discounting timescales.

These four phases are altered by predictive coding, especially phase 2) where the ramping upward is preceded by the adaptation from the cue-triggered jump and phase 3) where the return to baseline is accompanied by an undershoot (blue line in Figure 3A). Multiple research groups have shown that serotonin neurons are transiently activated by reward-predicting cues *in vivo* (11–13; *e*.*g*., Figure 3B inset; schematized in Figure 1A). Previous value and reward-based serotonergic theories cannot explain this phasic activity (black line in Figure 3A), leading to proposals that serotonin might encode some other quantity (*e*.*g*. surprise 13). In value prediction, phasic cue-associated firing emerges naturally as a result of adaptation (blue line in Figure 3A). Unlike previous models, our theory further predicts a subtle, counter-intuitive drop in activity during the reward epoch to complement phasic activation by the cue. Interestingly, this phenomenon is visible in raw experimental data presented in the literature (11–13), but is generally not quantified. In short, value prediction through adaptation explains why serotonin neurons are phasically activated by reward-predicting cues (Figure 1A) and predicts that serotonin neurons should exhibit decreasing/below baseline activity during reward consumption.

### Value prediction captures response to punishments

A significant problem for value and reward-based serotonergic theories is that serotonin neurons are often activated by both rewards and punishments (5, 11, 13). Since punishments can be seen as negative rewards *R*_*t*_ *<* 0, and value represents an estimate of future reward (Figure 2D), then serotonin neurons should be inhibited by punishments — not activated — if they encode a simple value signal (black line in Figure 3B).

In contrast, activation by punishments is expected under the value prediction theory. This is because predictive coding through adaptation (Figure 2G) creates an overshoot in the level of activity as the punishment ends (blue line in Figure 3B). To understand why this happens, recall that the effect of predictive coding through adaptation is to exaggerate positive (and negative) transients in the underlying value signal. The value signal during a punishment trial is the mirror image of the value during a reward trial (black lines in Figure 3A and B), increasing as the punishment ends just as the reward trial value signal decreases when the reward is consumed (3 s to 4 s post-cue in Figure 3A and B). Through predictive coding, the fast increase in value during the punishment epoch is enhanced, causing the encoded signal to briefly overshoot its baseline (*∼*4 s post-cue in Figure 3B).

The idea that predictively encoding a value signal creates a punishment withdrawal-induced overshoot explains 1) why serotonin neurons are activated by punishments as well as rewards (5, 11, 13), 2) why these seemingly opposite response features are positively correlated across cells (11), and 3) why this activation occurs at the end of a punishment rather than the beginning (4).

### Tonic firing during inter-trial intervals reflects reward in future trials

The phasic responses of serotonin neurons to rewards and punishments (Figure 1A and D) have historically been difficult to reconcile with tonic activity that tracks reward and punishment context (Figure 1B; 11, 35), spawning proposals that serotonin neurons may multiplex unrelated quantities over short intra-trial and long inter-trial timescales (11, 15). Value prediction explains these phasic responses (see above) while also predicting that tonic activity should track reward and punishment context (Figure 3C, Appendix E), unifying the responses of serotonin neurons to rewards and punishments over short and long timescales.

More interestingly, our theory predicts that trial duration should have pronounced effects on both inter-trial value coding and within-trial activity dynamics of serotonin neurons. Analysis of our model shows that the proportionality between inter-trial and within-trial value depends on two factors: 1) the mean duration of the ITI relative to the trial duration (vertical axis in Figure 3D) and 2) the duration of the trial (defined as the time between cue onset and reward delivery) relative to the discounting timescale of the animal *τ* (horizontal axis in Figure 3D). However, while the effect of ITI duration is surprisingly weak in the typical experimental range (*i*.*e*., ITIs two to five times the trial duration; 11–13), the effect of trial duration is quantitatively large and visually obvious. Specifically, when the trial duration is shorter than the discounting timescale, we expect to see both phasic cue-associated activity (Figure 3D1 and 3) and inter-trial value coding, while when the trial duration is longer than the discounting timescale, we expect ramping within-trial activity (Figure 3D2 and 4) and little to no inter-trial value coding. The transition between these two regimes is sharp and occurs when the trial duration is roughly equal to the discounting timescale. Thus, the ratio between the trial duration and discounting timescale controls both inter-trial value coding and within-trial “peak and plateau” vs. ramping activity dynamics.

**Fig. 4.**
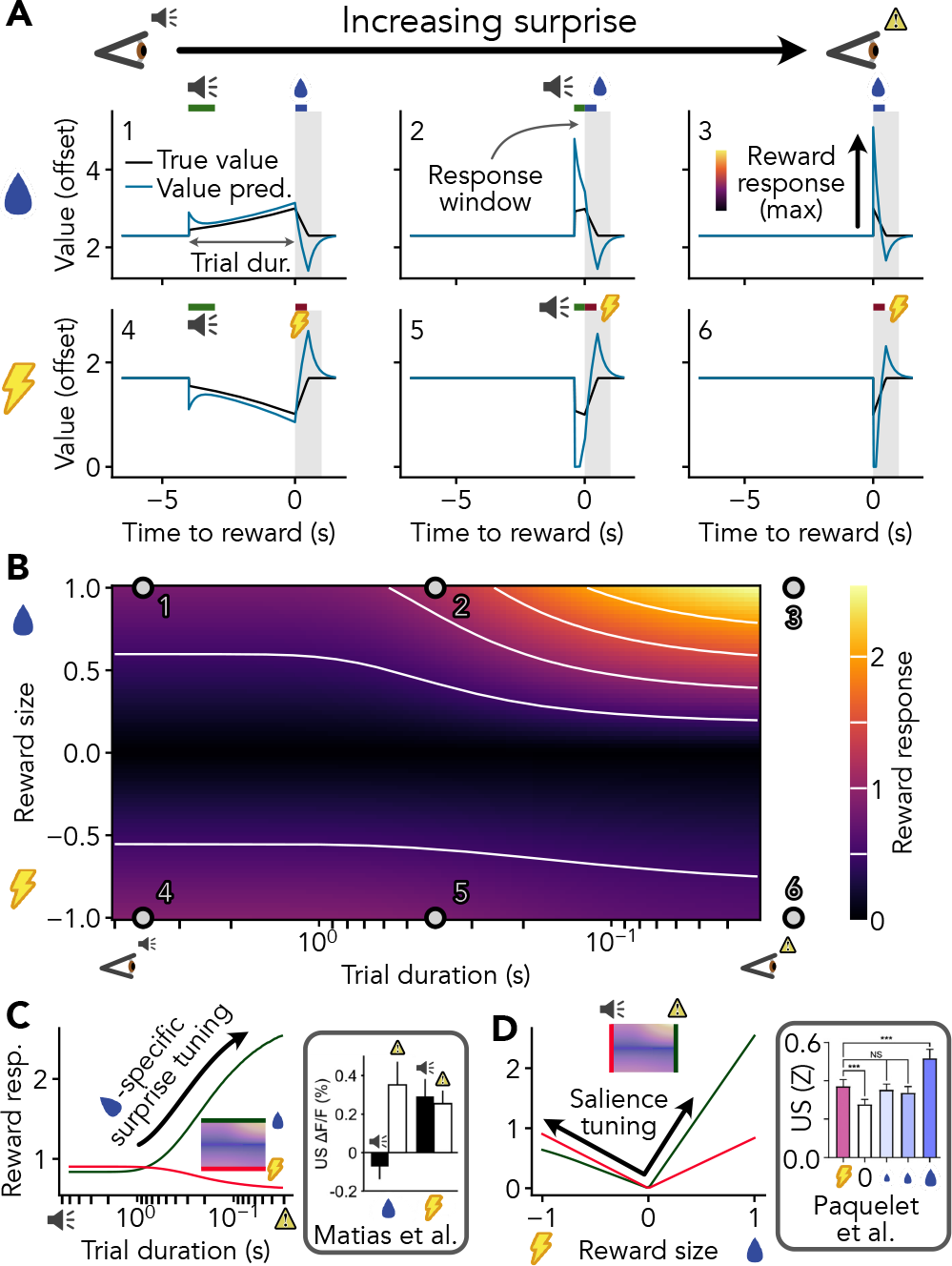
Predictively-encoded value resembles surprise and salience. **A**,**B** Reward responses depend on reward size and cue timing. Reward response is defined as the baseline-subtracted maximum DRN-encoded value signal within 1 s of cue onset (gray window). Trial duration is defined as the time between the onset of cue and reward (gray arrow in 1). Sample traces 3 and 6 represent uncued rewards and punishments, respectively. Signals are shifted up by 2 AU to capture decreased activity during punishment trials. **C** Predictively-encoded value yields larger reward responses for uncued vs. cued rewards but similar responses to uncued and cued punishments. Note resemblance to reward-specific surprise coding from Matias *et al*. (13) (inset modified from ref. Fig. 7B; punishment type: air puff). **D** Apparent salience coding is distinct from surprise. Whether cued or uncued, reward responses extracted from predictively-encoded value signals are smaller for neutral outcomes than both rewards and punishments. Note resemblance to salience coding from Paquelet *et al*. (5) (inset modified from ref. Fig. 2D with kind permission from Bradley Miller; punishment type: bitter quinine solution, reward type: sucrose solutions). Green and red lines in B and C are slices of data from A as indicated on mini-heatmaps.

The effect of trial duration on within-trial activity dynamics and inter-trial value coding predicted by our model explains 1) why “peak and plateau” dynamics and tonic value coding co-occur (11), 2) why experiments using longer trials sometimes produce ramping rather than “peak and plateau” dynamics (compare 11 and 12; schematized in Figure 1A; but see 13), and 3) why firing during the prereward epoch may decrease slightly as the trace duration increases (36).

### Value prediction explains reward-specific surprise

While little is known about the effect of trial duration on the pre-reward activity dynamics of serotonin neurons, the effect of trial duration on the amplitude of the reward (or punishment) response itself has received more attention. Compared with rewards delivered at the end of a trace conditioning trial, serotonin neurons are more strongly activated by rewards delivered spontaneously (11) or immediately following a cue (13). This has been interpreted as evidence for surprise coding (13), defined as activity that reflects an unsigned reward prediction error |*δ*_*t*_| = |*R*_*t*_ *−v*_*t*_|, which is believed to be important for learning (13, 15, 37). However, because the surprise/absolute RPE-like reward responses of serotonin neurons do not evolve on the same timescales as dopaminergic RPEs *δ*_*t*_ (13), are smaller than corresponding dopaminergic responses (11), and seem to be specific to rewards (Figure 1C and D; 11, 13), a different explanation is needed.

To understand how surprise tuning for rewards might emerge, we simulated value prediction under progressively shorter trace conditioning trials. As the trial duration shortened, the adaptation-induced cue-associated peak began to overlap with the response to the reward itself (compare Figure 4A1 and 2). This phenomenon becomes increasingly pronounced as the trial duration falls below the effective timescale of adaptation (on the order of hundreds of milliseconds; scan from left to right along the top of Figure 4B corresponding to the green line in Figure 4C), and is maximally strong when the trial duration reaches the zero lower bound, corresponding to an uncued reward (Figure 4A3).

It is difficult to differentiate absolute RPE from value prediction on the basis of reward responses alone because both theories predict stronger responses for surprising rewards (schematized in Figure 1C). To rule out absolute RPE, we turn our attention to serotonergic responses to punishments. Whereas the absolute RPE theory predicts that serotonin neurons should respond most strongly to surprising punishments (13), just as they do for rewards, serotonin neurons should have a very slight preference for predicted punishments under value prediction (Figure 4A4–6, left to right along the bottom of Figure 4B, red line in Figure 4C), consistent with experimental observations (13; schematized in Figure 1D).

To understand why value prediction implies reward-specific surprise tuning, recall that punishment-associated activity is caused by punishment *withdrawal* under our theory (Figure 3B). Shortening the trial duration has no effect on the rate of punishment withdrawal, and even causes the pre-punishment inhibition to slightly overlap the withdrawal-associated peak if the trial duration is sufficiently short, leading to a small decrease in the punishment response (Figure 4A4–6). The transition from surprise tuning to lack thereof occurs sharply when the size of the reward passes below zero, but this is obscured by the relatively small responses to near-zero rewards in our model (Figure 4B). As with rewards, the transition to a slight preference for unsurprising punishments occurs when the trial duration drops below the effective timescale of adaptation, such that reward and punishment responses are expected to diverge markedly for trial durations on the order of hundreds of milliseconds or less (Figure 4C).

These simulations show that value prediction explains surprise tuning for rewards that reverses for punishments, thus providing a more complete account of serotonergic surprise tuning than the existing absolute RPE theory (13).

### Value prediction explains salience tuning for both surprising and unsurprising stimuli

If surprise is defined in the serotonin literature as absolute RPE |*δ*_*t*_|, then salience is defined as the absolute size of the reward itself |*R*_*t*_|. We have already shown that value prediction explains serotonergic responses to cued rewards and punishments (Figure 3A and B), a type of salience tuning, but recent experimental work has focused on this phenomenon in the context of *uncued* rewards and punishments (5).

To show that value prediction explains salience tuning for both uncued and cued rewards, we simulated trace conditioning experiments using a wide range of trial durations and reward sizes. We find stronger responses to both rewards and punishments compared with neutral outcomes across all trial durations, consistent with salience tuning (Figure 4B). Interestingly, the salience tuning effect is fairly balanced for cued rewards and punishments (“V”-shaped red line in Figure 4D), whereas a clear preference for rewards emerges in very short trials (“✓”-shaped green line in Figure 4D). This reward preference can be explained by the interaction of reward-specific surprise tuning (Figure 4C) with salience. Thus, evidence for existing salience and surprise theories also supports value prediction.

### Slow online learning

So far we have focused on the resemblance between predictively-encoded true value signals and the *in vivo* activity patterns of serotonin neurons. However, it is unrealistic to think that serotonin neurons signal true value, since this would require perfect knowledge of future rewards (Figure 2D). Instead, serotonin neurons likely encode a value signal that is estimated on the basis of past rewards (Figure 2E).

Does our focus on true rather than estimated value pose a problem for the results presented above? To find out, we applied an online value estimation algorithm (van Seijen’s TD(*λ*) [40], see Methods) to a trace conditioning experiment consisting of hundreds of trials. The estimated value signal exhibited a ramp that gradually increased in amplitude, gradually converging to a close approximation of the true value (Figure 5A), mirroring observed activity of serotonin neurons in mice (12, 13). The same was true of the predictively-encoded estimated value (Figure 5B). Overall, estimated value signals resembled a true value template rescaled by reward history. These simulations illustrate that the details of how the value signal is calculated (*i*.*e*. on the basis of future rewards, as in true value Figure 2D, or on the basis of past rewards, as in estimated value Figure 2E) play only a minor role in shaping the activity patterns of serotonin neurons during trace conditioning, whereas predictive coding and reward history are critical.

**Fig. 5.**
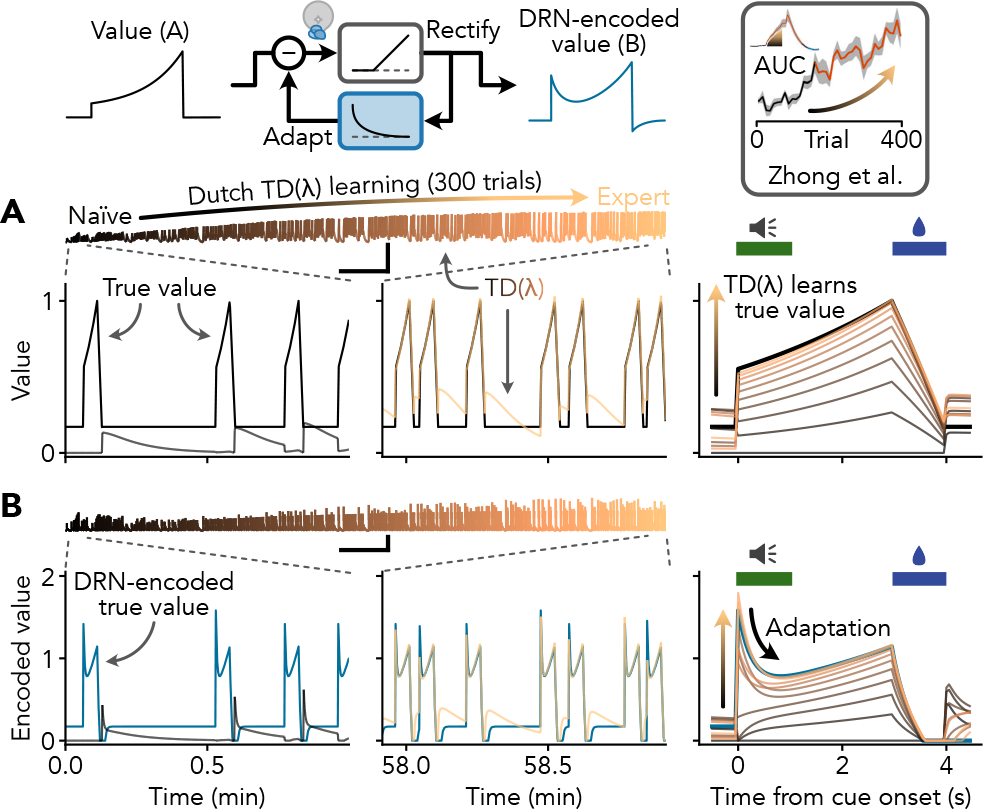
Online value estimation. **A** Estimation of value during 300 trials of trace conditioning. Comparison between true value *v*_*t*_ (black) and value estimated using TD(*λ*) with Dutch eligibility traces (shades of copper). Note that the estimated value signal converges to a close approximation of the true value (right). **B** Value signals predictively encoded by the DRN. Scale bars: 5 min, 1 arbitrary value unit. Vignette above A modified from ref. Fig. 1G of Zhong *et al*. (12).

If the activity levels of serotonin neurons are scaled by reward history, how large should this effect be? The rate at which the TD(*λ*)-estimated value signal is scaled up and down by reward history depends on the learning rate *α*, which can also be expressed as the estimation timescale *τ*_est_ = *−dt* ln(1 *− α*). Previous work suggests that the timescale over which serotonin neurons integrate rewards is on the order of hundreds of trials (12, 13). Since the estimation timescale is so long, it is possible to write a first-order approximation of the effect of an uninterrupted string of rewards (or reward omissions) on the firing rates of serotonin neurons (Appendix G. Using an estimation timescale of *τ*_est_ = 200 trials (13), background firing rate of 2 Hz (11, 15), and winning streak of five trials, we expect to see firing rate modulations of only *∼*0.05 Hz. While longer runs of rewards (or reward omissions) would produce larger effects (*e*.*g*., 0.4 Hz for a run of 50 trials), such winning streaks (or losing streaks) are rare in experiments with probabilistic rewards that are best suited to studying the effects of reward history on serotonergic activity (15). In short, if serotonin neurons encode a value signal that is estimated over a long period of time, as evidence suggests (12, 13), we expect to see only small effects of reward history on serotonin neuron firing (*≪*1 Hz) in typical experiments.

### Serotonin neurons quantitatively encode reward history

Value prediction unifies a wide range of experimental observations that do not have a consistent interpretation under existing serotonergic theories (Figure 1, Table 1). To assess whether value prediction generalizes beyond the main qualitative results of Cohen *et al*. (11), Zhong *et al*. (12), and Matias *et al*. (13), we re-analyzed a dataset of serotonergic responses to dynamically-varying *in vivo* rewards from Grossman *et al*. (15). To begin, we sought to determine whether the activity levels of serotonin neurons in this dataset are weakly modulated by reward history, and, in particular, whether this is true of neurons with activity dynamics resembling a predictively-encoded value signal.

A short description of the experiment and our analysis approach follows (see Methods for details). The dataset consists of tetrode recordings of *N* = 37 identified serotonin neurons in mice receiving dynamically-varying probabilistic rewards in a trace conditioning experiment (15, 23). We extracted the spikes of each neuron in a short window around each trial along with the proportion of the past five trials that were rewarded (Figure 6A). Since true value and TD value estimates are both very closely tied to the mean reward (Appendix B), we use the five-trial mean reward as an interpretable proxy for value. Because the effect of reward history on serotonin neuron activity is expected to be small (see previous section) and many common statistical tests for value coding are prone to high rates of false positives (41), we used circular trial permutation (Figure 6B). This test breaks the putative association between reward history and serotonin neuron activity while preserving all other structure in the data (for example, slow fluctuations related to arousal). If the association between serotonin neuron activity and reward history is stronger when the trials are correctly aligned than when this alignment is broken, we can conclude that the correlation between serotonin neuron activity and reward history cannot easily be explained by random fluctuations in the data.

**Fig. 6.**
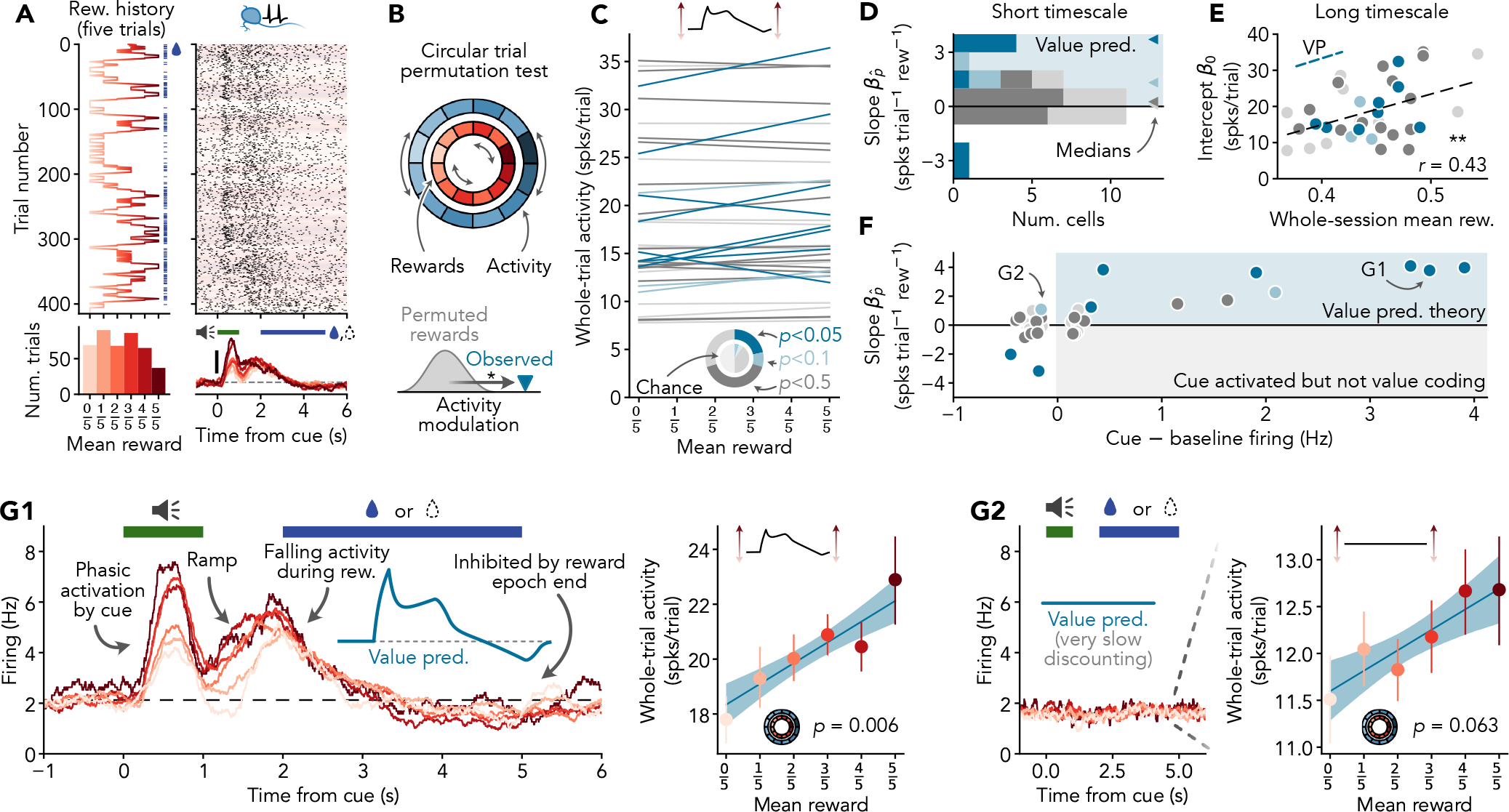
Individual serotonin neurons exhibit reward coding features consistent with value prediction. **A** Example trace conditioning experiment of Grossman *et al*. (15) (see Methods). Counterclockwise from top left: Bernoulli water rewards (blue lines) and time varying reward probability estimated using a five-trial moving average (red line). Distribution of number of trials at each level of estimated reward probability; note the wide range of reward probabilities in this dataset. Firing rate of the serotonin neuron recorded in this session calculated using a 500 ms PSTH, lines coloured according to estimated reward probability as in histogram at left, scale bar 2 Hz. Spike raster used to calculate PSTH, background is shaded according to the estimated reward probability as in the other plots. **B** Circular trial permutation test used to assess statistical significance of correlations between estimated reward probability and serotonin neuron activity. Reward history is randomly shifted with respect to spiking activity to build up a null distribution against which the observed correlation can be compared. **C** Serotonin neuron whole-trial activity reflects reward history. Each line represents the following regression model fitted to a single neuron: 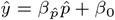, where 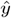 is the predicted whole-trial activity (defined as the number of spikes within a 7.5 s period beginning 1.5 s before the start of the cue and ending 1 s after the end of the reward epoch), 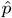 is the reward probability estimated as in A, and 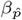 and *β*_0_ are the slope and intercept. Slope 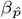 represents the effect of recent reward history 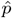 on activity and intercept *β*_0_ represents the baseline activity level following a short string of unrewarded trials 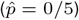. Donut plot shows the distribution of circular trial permutation test *p*-values against 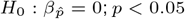 occurs significantly more frequently than 5 % chance rate (inner pie chart). Lines are colour-coded according to statistical significance of the slope 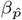 as in the donut plot. *N* = 37 neurons. **D** Distribution of regression slopes 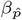. Note tendency towards positive slopes consistent with value prediction. Colour-coded as in C. **E** Distribution of regression intercepts *β*_0_ . Note positive correlation between baseline activity and reward rate over very long timescales consistent with value prediction. Colour-coded as in C. **F** Relationship between reward history modulation (vertical axis) and phasic cue-associated firing (horizontal axis). Note that neurons with clear cue-associated firing (G1 and A, for example) tend to be positively modulated by reward history, consistent with value prediction. **G** Firing dynamics and whole-trial activity modulation in representative serotonin neurons. G1 shows a neuron with clear trial-associated activity dynamics and positive activity modulation by reward history, representative of neurons in the upper right of F. Note the striking correspondence between activity dynamics of the value prediction model (blue inset) and the example neuron. G2 shows a neuron with no clear trial-associated activity dynamics and numerically positive (but not statistically significant) activity modulation by reward history, representative of most neurons in the left part of F. Regression plots (G1 right and G2 right) illustrate the analysis used for C–F. Error bars/bands represent 95 % bootstrap confidence intervals (with Monte-Carlo bias correction in the case of error bars) provided for illustration purposes only.

With data and statistical approach in hand, we turned our attention to quantifying the effect of reward history on the activity levels of serotonin neurons. Regressing the number of spikes per trial (which we refer to as whole-trial activity) onto the five trial mean reward for each neuron (Figure 6C) revealed statistically-significant modulation by reward history in 8*/*37 cells (circular permutation test *p <* 0.05 against regression slope 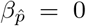; significantly above the 5 % chance rate, binomial proportion test *p <* 0.001; see donut plot in Figure 6C) somewhat variable levels of background activity (regression intercepts *β*_0_ = 18.6 *±* 8.0 spikes trial^*−*1^, mean *±* SD, equivalent to 2.5 *±* 1.1 Hz, coefficient of variation 0.43). It is possible that 22 % represents a lower bound on the proportion of cells in our sample that encode value. Consistent with this possibility, the relationship between activity and reward history was generally positive across the population (regression slopes 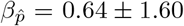 spikes trial^*−*1^ reward^*−*1^, Wilcoxon signed rank test *p* = 0.039; equivalent to 0.09 *±* 0.21 Hz, roughly consistent with the effect size calculation in the previous section; 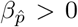 in 65 % of cells, one-sided sign test *p* = 0.049), including in many cells that did not cross the *p* = 0.05 significance threshold in the circular trial permutation test (Figure 6D, note significance-stratified medians). We observed two neurons with a statistically significant negative effect of reward history on whole trial activity (Figure 6C and D). This is consistent with the expected false positive rate (2*/*37 = 5.4 %), but it is also possible that value prediction is not universal in the DRN. Finally, we also observed a significant correlation between background activity and the proportion of all trials rewarded in the corresponding session (Pearson *r* = 0.43 between regression intercept *β*_0_ and whole-session mean reward, *p* = 0.008, Figure 6E), consistent with value coding over timescales beyond the five trial horizon. This correlation is surprisingly strong considering the many factors that impact background firing rate (differences in the biophysical features or inputs of individual serotonin neurons, differences in thirst or arousal between recording sessions and mice, etc.) and the coarseness of the whole-session reward metric. Due to a high rate of statistically null results which neither confirm nor rule out value coding, additional analyses are needed to determine to what extent value prediction is typical in the DRN (see “Value prediction dominates population activity” below). For now, we conclude that at least some serotonin neurons exhibit positive reward history modulation consistent with value prediction.

Our theory predicts that the phasic reward cue-associated activity observed in some serotonin neurons is due to value coding, while other theories propose that phasic activity could be unrelated to reward history. To examine the potential connection between this phasic activity and value prediction, we stratified the slopes obtained from our regression analysis according to the amplitude of the cue-associated extremum in the firing rate (Figure 6F). Of the small number of cells with a clear cue-associated peak (*N* = 7 cells with *>*1 Hz increase in firing above baseline), all exhibited numerically positive reward history modulation (one-sided sign test *p* = 0.008, *N* = 7) which was individually statistically significant in just over half of these cells (circular trial permutation test *p <* 0.05 in 4*/*7 neurons; exact *p* values for each neuron are 0.006, 0.007, 0.009, 0.013, 0.071, 0.126, 0.237 in order of decreasing significance, values lower than *p* = 0.002 are not possible). The connection between phasic activity and value coding was surprisingly consistent: out of 37 neurons, we did not observe any with both clear phasic activity and numerically negative reward history modulation (bottom right quadrant in Figure 6F). We conclude that there is a strong association between phasic cue-associated activity and positive modulation by reward history, consistent with the idea that value prediction underlies both phenomena.

The main ideas of this analysis are summed up by the two example neurons presented in Figure 6G. In Figure 6G1, we see a neuron with activity dynamics strikingly similar to the value prediction model, including cue-associated phasic firing, a pre-reward ramp, and falling activity during the reward period. In this neuron, we also observe statistically significant (circular trial permutation test *p* = 0.006) and almost perfectly linear positive scaling of whole-trial activity by reward history, consistent with value prediction. The neuron in Figure 6G1 is clearly well-described by our theory, but the same cannot be said of the neuron in Figure 6G2. Examining its peri-stimulus time histogram (PSTH) reveals no clear activity dynamics, and there is no statistically-significant effect of reward history on firing (circular trial permutation test *p* = 0.063). This neuron may not predictively encode value, but it is also possible that the timescale of this experiment is simply too fast. An absence of discernible within-trial activity dynamics is consistent with value prediction if the discounting timescale is much longer than the trial duration^1^ (Figure 8), and, while not statistically significant, the magnitude of the reward history modulation is actually closer to our predictions than the unusually large effect shown in Figure 6G1 (for an increase in reward probability from 1*/*2 to 1, the calculation in the previous section shows we expect an increase in firing of *∼*0.05 Hz, compared with 0.25 Hz and 0.07 Hz in example neurons 1 and 2, respectively). The analyses presented in this section prioritize clarity over statistical power. As a result, effects that pass the statistical significance threshold are likely to be unusually strong and not representative of the broader population of value coding serotonin neurons (42). Our results are best interpreted as a lower bound on the proportion of serotonin neurons that encode value.

**Fig. 7.**
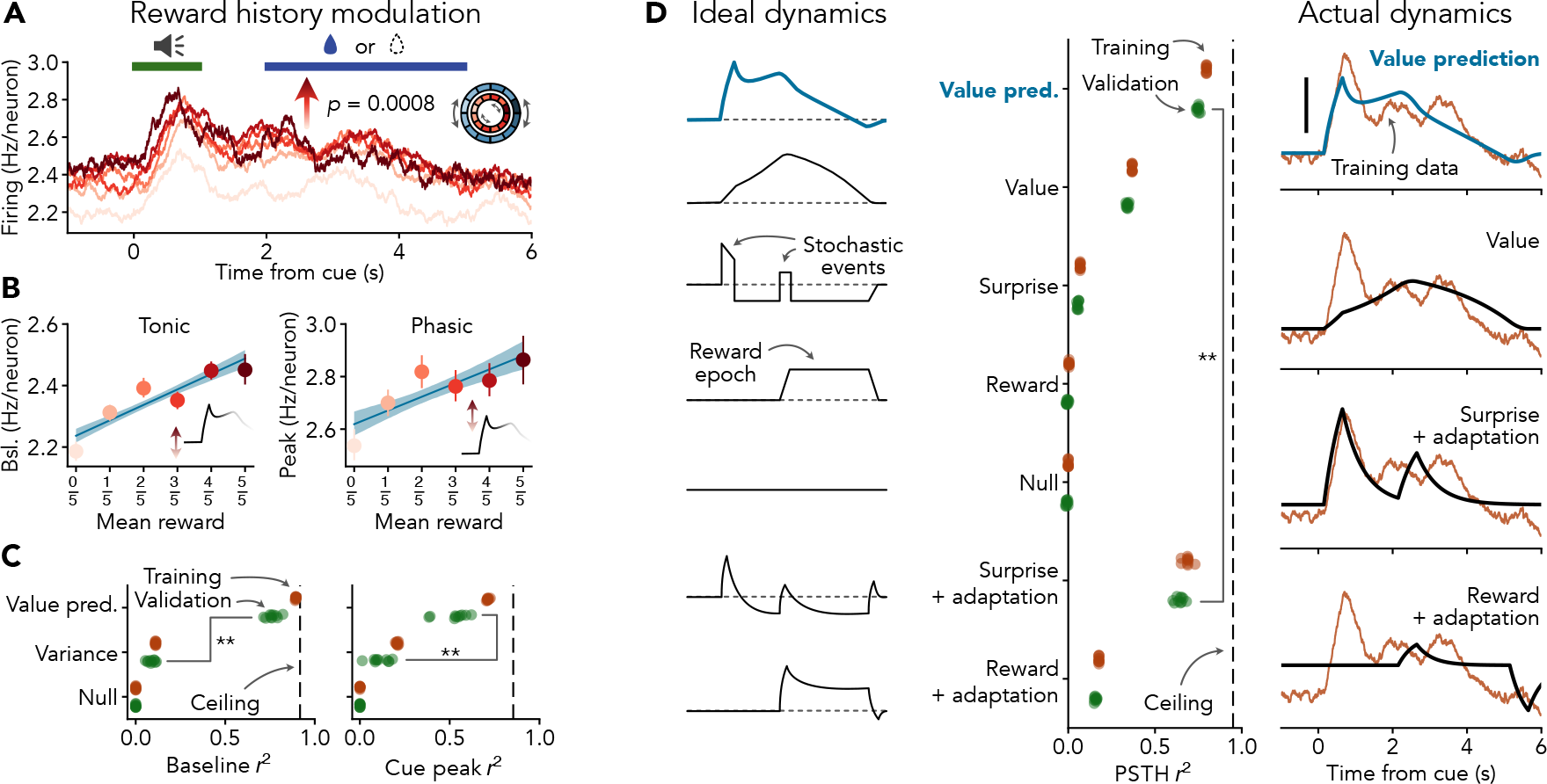
Value prediction better explains serotonin neuron population activity than competing theories. **A** Whole-trial population activity encodes reward history. Whole-trial activity and circular permutation test as in Fig. 6. **B** Baseline population firing rate quantitatively encodes reward history. Baseline activity defined as mean of PSTH 1 s before start of cue. Peak cue activity is defined as the maximum of the PSTH during the 1 s cue period. Error bars represent 95 % confidence intervals obtained via bootstrap with Monte-Carlo bias correction. Error bands around regression lines represent 95 % confidence interval obtained via bootstrap. Regression slopes are significantly different from zero; bootstrap 99 % CI test. **C** Value prediction better accounts for reward history modulation of population firing rate than variance. *x*-axis represents the proportion of variance explained (weighted *r*^2^) by each model fitted to data as in B. Performance is presented as the mean five-fold cross-validated accuracy, each point represents one cross-validation repeat. Ceiling line represents the accuracy obtained using the training data to predict the validation data directly (maximum across all repeats). **D** Value prediction better accounts for population firing rate dynamics than competing theories. Schematics at left illustrate predictions of each theory; note that surprise-like signals should decrease during trials, but this does not happen in our model fits. Performance is assessed using repeated five-fold cross-validation as in C. Scale bar 0.2 Hz neuron^*−*1^. Trial structure as in A.

**Fig. 8.**
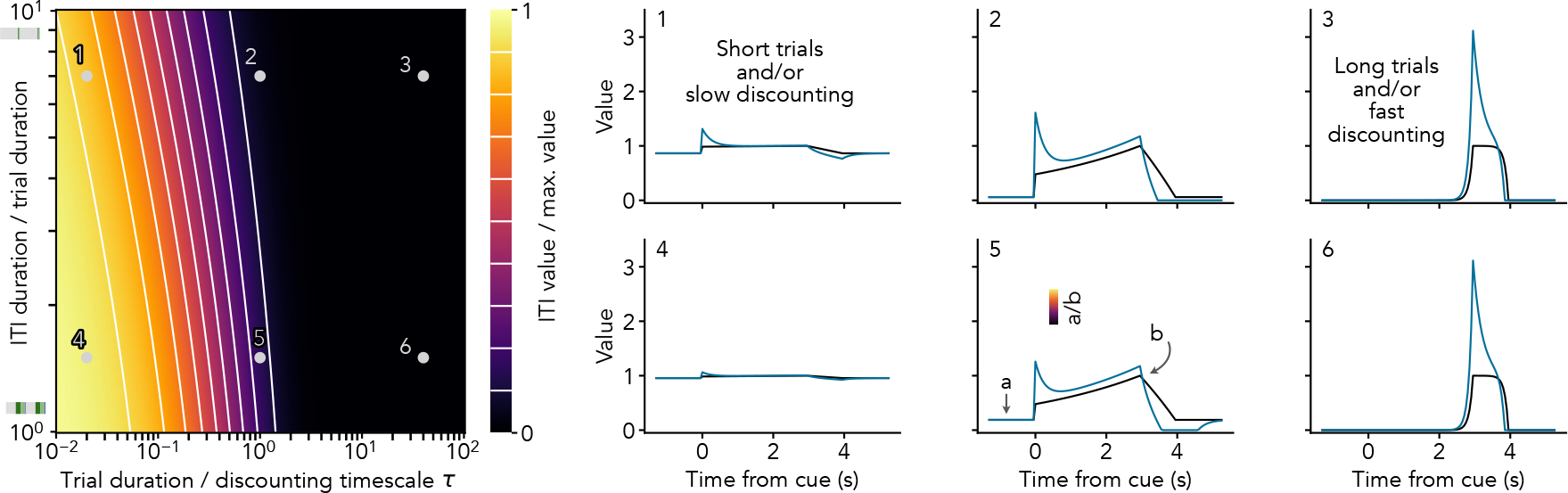
Effects of discounting over extreme timescales. This figure extends the horizontal axis of main text Figure 3D. Heatmap shows the ITI value relative to the maximal value during the trial (*i*.*e*., the value just before the start of reward delivery) as a function of experimental parameters (vertical axis) and the duration of the trial relative to the discounting timescale of the agent (horizontal axis). Ribbons next to the vertical axis are to scale, gray represents the mean ITI duration and colours represent trial epochs. Traces show the true value dynamics (black) and predictively-encoded value (blue) for a trial consisting of a 3 s combined cue and delay epoch and a 1 s reward epoch. True value is normalized to the maximum just before reward as in the heatmap; different reward sizes (and learning) can be accommodated by scaling the traces. Note that the value signal is nearly flat if the discounting timescale is much longer than the trial duration (traces 1 and 4), similar to the dynamics of some serotonin neurons (*e*.*g*., Figure 6G2).

In this section, we have shown that value prediction provides a very good description of the activity patterns of at least some serotonin neurons. Could a different model provide an equally good or even better description of serotonin neuron activity? To what extent are the tuning features explained by value prediction dominant at the population level? We turn to these questions next.

### Comparison with expected uncertainty

A significant body of literature argues that serotonin neurons encode a quantity related to RPE, typically its absolute value, in order to signal surprising events (7, 13, 15). Perhaps surprisingly, some of the strongest evidence for this perspective is consistent with our results.

An influential model of the role of serotonin in learning connects trial-to-trial modulations of serotonin neuron activity to a moving average of absolute RPEs called expected uncertainty (15). To compare expected uncertainty against our model (which predicts an essentially linear relationship between mean reward and serotonergic activity, *e*.*g*. Figure 6G1 right), we analytically derived the relationship between reward probability and mean absolute RPE in models of animal learning (Appendix F). While the mean absolute RPE is precisely twice the variance of a binary reward in the simplest of RL models (and therefore has an inverted U-shaped relationship with reward probability), sophisticated models of animal learning often include features that profoundly alter this relationship (*e*.*g*., 15, 18). As a result, in theory expected uncertainty is usually negatively related to the mean reward in addition to being positively related to variance (Figure 11). Re-analyzing the computed expected uncertainty values from the dynamic Pavlovian task in Grossman *et al*. (15) shows that this is also true in practice: expected uncertainty is more strongly correlated with reward probability than variance in 26*/*28 sessions in this dataset (Figures 9 and 10; median marginal *r*^2^ between expected uncertainty and five-trial mean reward 0.815, IQR 0.565 to 0.903, compared with median 0.083, IQR 0.016 to 0.216 for variance). The fact that the previously-reported correlation between expected uncertainty and serotonergic activity is negative more often than not also suggests a connection between serotonergic activity and mean reward rather than variance. We conclude that evidence for serotonin neurons signalling expected uncertainty is consistent with value coding.

**Fig. 9.**
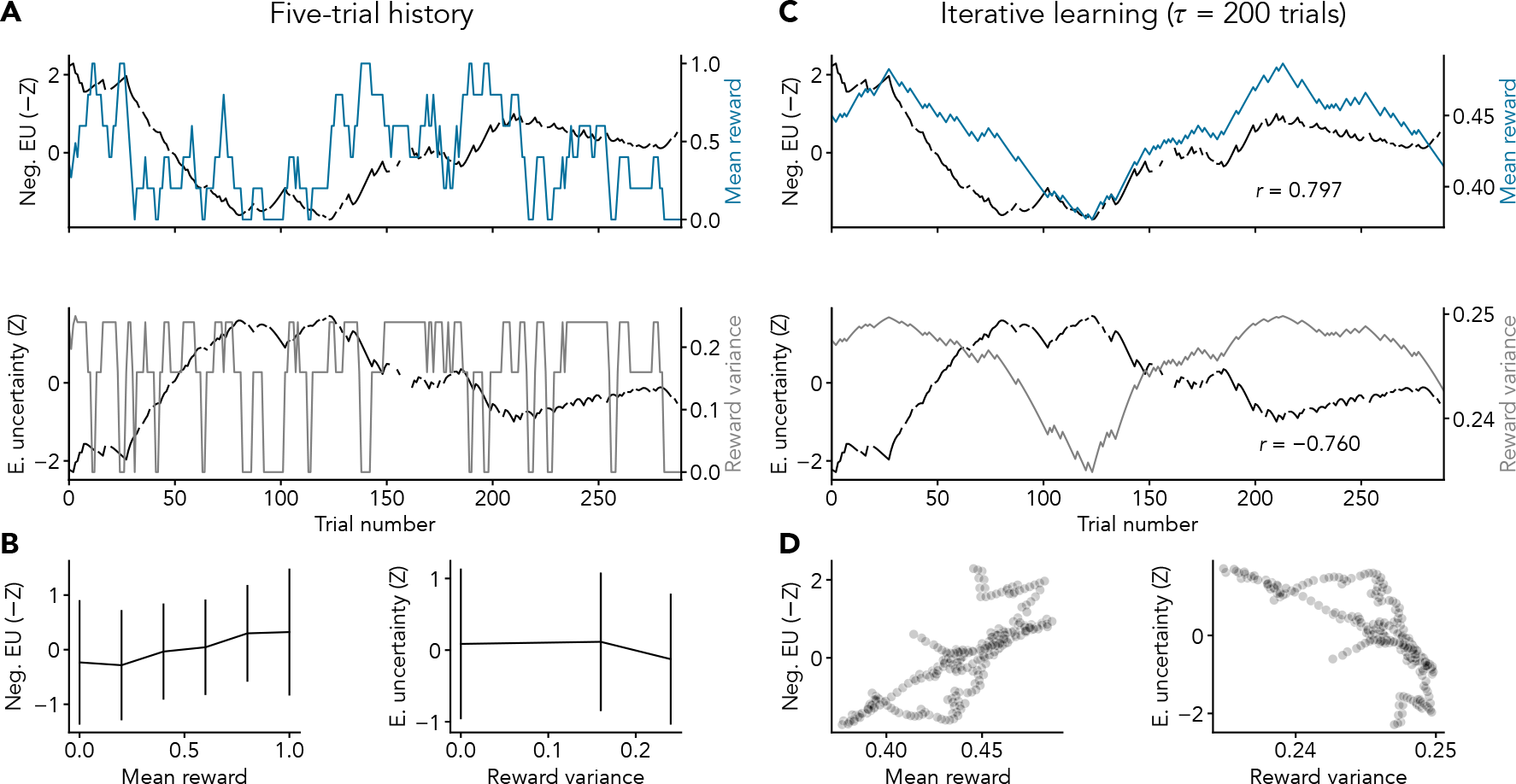
Expected uncertainty (15) reflects mean reward, not reward variance, in an example session. **A** Comparison between the time-course of expected uncertainty (black) and mean reward (blue, top) or reward variance (gray, bottom) calculated using a five-trial history. Note that expected uncertainty is theoretically *negatively* related to the mean reward and *positively* related to variance (Appendix F). Consistent with this, negative expected uncertainty closely resembles five-trial mean reward (top). Gaps in the expected uncertainty timeseries are due to removal of catch trials. **B** Marginal relationship between expected uncertainty and mean reward (left) and reward variance (right) calculated using a five-trial history, as in our main analysis (*e*.*g*., Figure 7B). Data are presented as mean *±* SD. Note connection between negative expected uncertainty and the mean reward (left). **C** Comparison between the time-course of expected uncertainty (black) and mean reward (blue, top) or reward variance (gray, bottom) calculated using an exponential moving average with an estimation time constant of *τ*_est_ = 200 trials. The mean reward calculated using an exponential moving average is equivalent to state value under TD learning (Appendix B). **D** Marginal relationship between expected uncertainty and mean reward (left) and reward variance (right) calculated using an exponential moving average as in C. Note connection between expected uncertainty and mean reward (left). Z-scored expected uncertainty values for recording session mBB036d20160623 (23) used in (15) kindly provided by Cooper Grossman.

**Fig. 10.**
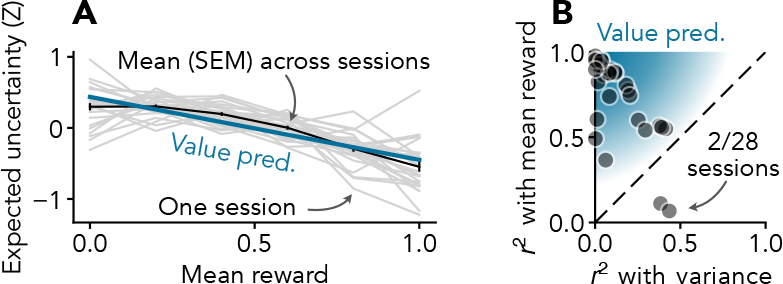
Expected uncertainty (15) reflects mean reward, not reward variance, across sessions. **A** Marginal relationship between expected uncertainty and mean reward. Marginals were calculated by averaging expected uncertainty at each level of mean reward calculated using a five-trial history, as in our main analysis (*e*.*g*., Figure 6A). Black line represents mean *±* SEM across *N* = 28 recording sessions. Note that expected uncertainty is negatively related to the mean reward, consistent with our simplified model (Appendix F). The relationship between expected uncertainty and five-trial mean reward is nearly linear, possibly due to the long timescale over which expected uncertainty is estimated (15). **B** Expected uncertainty is more closely related to the mean reward than reward variance in nearly all sessions. Unweighted *r*^2^ values represent the fraction of variance in the gray lines from A that is explained by a straight line (mean reward, vertical axis) or a parabola (variance, horizontal axis). Z-scored expected uncertainty values used in (15) kindly provided by Cooper Grossman.

**Fig. 11.**
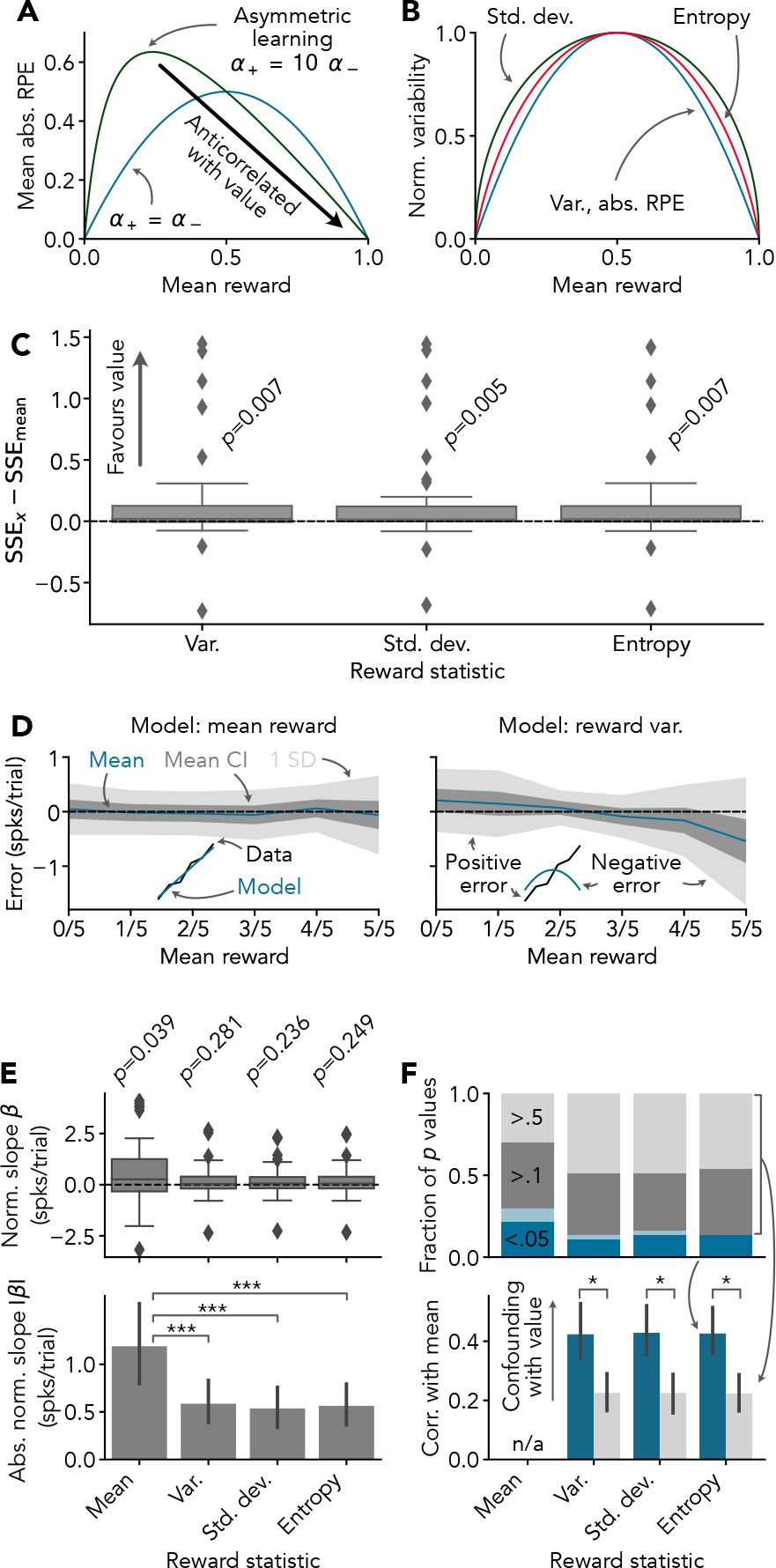
Serotonin neuron activity reflects mean reward and is spuriously correlated with reward variability. **A**,**B** Reward variability is inextricably linked to mean reward for binary (Bernoulli) rewards. **A** Expected uncertainty (15), defined as the mean RPE in a model of biased animal learning, has an inverted U-shaped relationship with the mean reward. If learning rates for positive and negative RPEs are highly asymmetric, as in previous work, expected uncertainty is mostly negatively related to the mean reward. **B** Unbiased statistics of reward variability, such as reward variance, also have inverted U-shaped relationships with the mean reward. **C** Serotonin neuron activity is better described by mean reward than reward variability. SSE*x −* SSE_mean_ represents the difference in weighted sum of squared errors (SSE) between a linear fit of serotonin neuron whole-trial activity to the reward variance, standard deviation, or entropy versus a linear fit of whole-trial activity to the mean reward (estimated using a five trial history, as in main text). Positive values indicate a better fit (lower error) with the mean reward model. *p* values are from Wilcoxon signed-rank tests. **D** Serotonin neuron activity does not systematically deviate from the mean reward model (left). Errors are derived from linear fits used in C. Dark gray bands indicate 95 % CIs on the mean (*i*.*e*., mean *±*1.96 SEM). No systematic errors are expected if the model is essentially correct (illustrated in inset). Note that the variance model exhibits systematic errors (right), as expected if serotonin neuron activity encodes the mean reward (illustrated in inset). **E** Serotonin neuron activity is relatively strongly positively related to mean reward. Slopes are derived from fits used in C and normalized to the dynamic range of the corresponding reward statistic as in B (0–1 for mean reward, 0–1*/*4 for variance, 0–1*/*2 for standard deviation, and 0–1 for entropy). *p* values are from Wilcoxon signed-rank tests. **F** The activity of individual serotonin neurons is more often correlated with mean reward than reward variability, and significant correlations with variability are likely due to confounding. A significant proportion of serotonin neurons exhibit activity that is significantly correlated with each reward statistic (top; cyclic permutation test, see main text; dark blue, light blue, dark gray, and light gray indicate *p ≤* 0.05, 0.05 *< p ≤* 0.1, 0.1 *< p ≤* 0.5, and *p >* 0.5, respectively, as in main text). However, in the subset of neurons in which activity is significantly correlated with reward variability, reward variability is unusually strongly confounded with the mean reward (bottom). Stars indicate 0.01 *< p ≤* 0.05 (actual range: *p* = 0.024 to *p* = 0.037) in permutation-based two sample *t* tests (non-parametric tests could not be used because too few neurons had statistically-significant correlations with reward variability statistics). *N* = 37 neurons in all cases.

### Comparison with reward variance

To address the possibility that serotonergic activity might encode reward variability in a way that is not captured by expected uncertainty, we repeated the regression analyses described above (Figure 6) using reward variance in place of the mean reward (Figure 11). We found that the whole-trial activity levels of serotonin neurons are better described by mean reward than variance (Wilcoxon signed-rank test on weighted sum of squared errors from regression fits *p* = 0.007, *N* = 37), and the effects of mean reward are larger (absolute change in activity of 1.19 *±* 1.25 spikes trial^*−*1^ from a reward of zero to a reward of one and 0.59 *±* 0.69 spikes trial^*−*1^ from a variance of zero to the maximum variance of 0.25, both quantified using the absolute regression slope, Wilcoxon signed-rank *p* = 0.001), more consistent (regression slopes are typically positive for mean reward, Wilcoxon signed-rank *p* = 0.039, but symmetric around zero for variance, Wilcoxon signed-rank *p* = 0.281), and statistically significant twice as often (21.6 % and 10.8 % of cells with circular trial permutation test *p <* 0.05 against regression slope equal to zero for mean and variance, respectively; for comparison, we expect significant *p* values in up to 12 % of cells even if none actually encode variance, approx. 95 % CI on a proportion of 5 % with *N* = 37).

Because the mean and variance of binary rewards are directly related, we wondered whether the apparent correlations between reward variance and serotonin neuron activity might actually be due to value coding. Consistent with this idea, statistical confounding between mean reward and variance was unusually strong in the subset of cells with a significant correlation between variance and activity (Pearson *r* between variance and mean reward is higher in cells with circular trial permutation test *p <* 0.05 for the variance slope *β* = 0, permutation *t* test *p* = 0.039).

Our simple analysis does not reveal clear evidence that serotonin neurons encode an unbiased estimate of reward variance. To mitigate the possibility that these findings are sensitive to technical details of our approach, we repeated the above analyses using reward standard deviation and entropy as alternative definitions of variability, using an iterative/TD method to estimate reward statistics rather than a five-trial moving average, and quantifying activity using a pre-trial baseline rather than whole-trial activity (Figures 11 and 12). None of these variations affected our results.

**Fig. 12.**
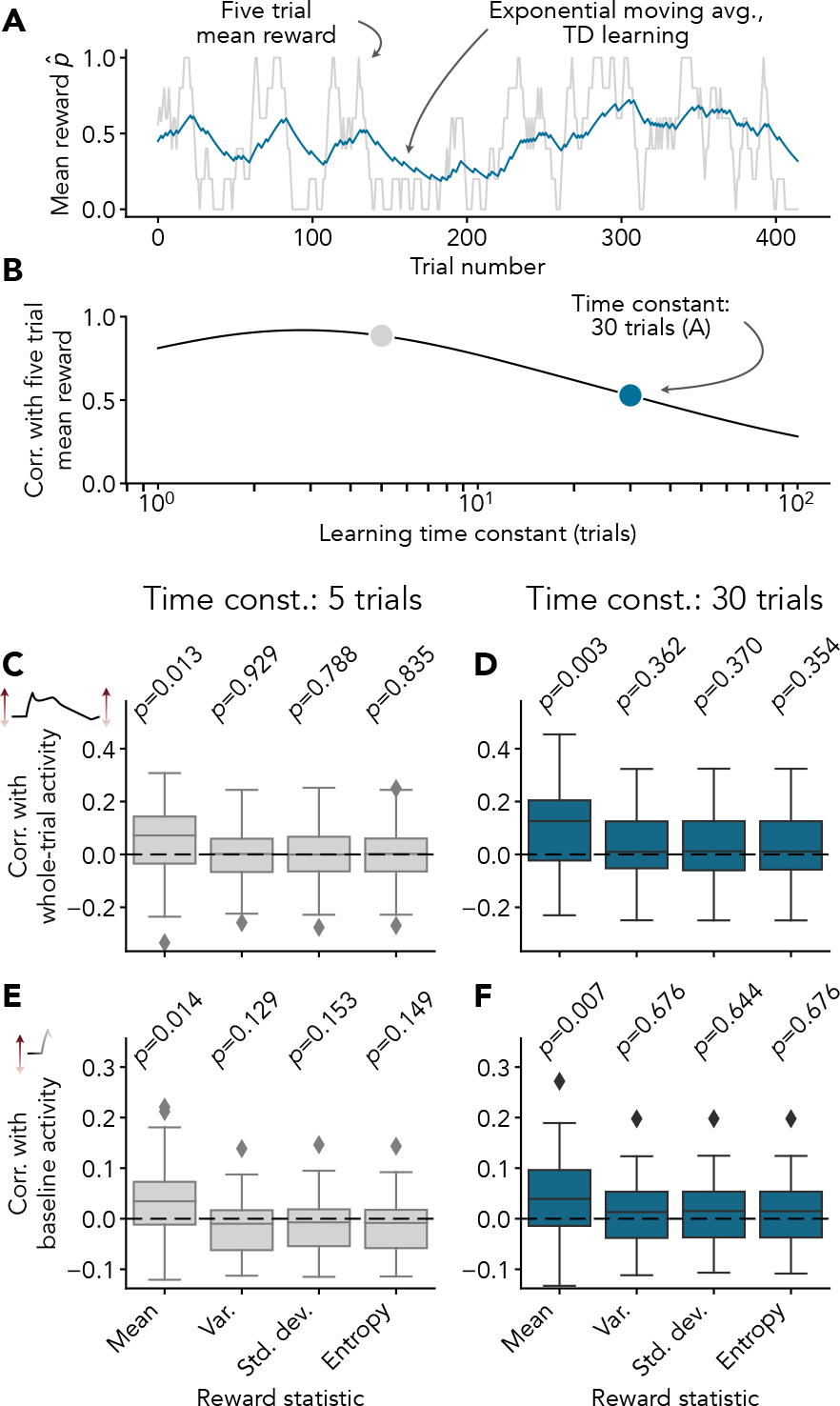
Correlation between serotonergic activity and mean reward does not depend on technical details of analysis. **A**,**B** Conclusions are unlikely to depend on the decision to use a five trial reward history as a proxy for value rather than an incrementally-learned estimate because the two are very similar. **A** Comparison of five trial mean reward used in main text with an exponential moving average of past reward for an example session. The blue line represents an exponential moving average with a time constant of 30 trials, which is equivalent to the value estimated by temporal difference (TD) learning using a learning rate of *α* = 1*/*30 (and no discounting). **B** Pearson correlation between five trial mean reward and moving reward average as a function of the time constant of the moving average (learning time constant) for the example session shown in A. Note that the two are moderately or strongly correlated across a wide range of learning time constants. **C**,**D** Whole trial activity is correlated with mean reward but not reward variability when reward statistics are calculated using a moving average of past rewards rather than a five trial history. Note that this result does not depend on the choice of learning time constant. **E, F** Choice of activity metric does not affect conclusions: baseline activity is also correlated with mean reward but not reward variability. In C–F, reward variance, standard deviation, and entropy were calculated from the learned 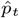 rather than moving averages (of squared errors, for example) because this approach is less statistically biased and/or better defined. *N* = 37 cells in all cases.

We are unable to conclude that serotonin neurons encode reward variability in this task. If serotonin neurons do encode reward variability, our results are consistent with a variability code that is significantly weaker and less consistent than the code for value.

### Untested models

To address the possibility that a different, untested model might provide a better account of the relationship between reward statistics and serotonergic activity, we inspected the residuals of our regression fits (Figure 11). Whereas the regression against reward variance systematically overestimates serotonin neuron activity when the mean reward is low and overestimates activity when the mean reward is high, we did not observe any obvious structure in the errors of the regression against the mean reward. The marginal relationship between serotonergic activity and mean reward is surprisingly well-described by a straight line, offering no hint as to what a better model might be.

### Value prediction dominates population activity

Value prediction provides a good description of the activity dynamics and reward history modulation of at least some serotonin neurons, but are the features captured by our model typical of serotonin neurons in general? To address this question, we constructed a synthetic serotonin neuron population using the *N* = 37 cells in our dataset and tested how well value prediction explains the synthetic population-level activity patterns in comparison with other models. If these cells do not generally predictively encode value, the features predicted by our model could be washed out by noise or masked by other coding features that are subtle at the single neuron level.

We began by considering whether serotonin neuron population activity is positively modulated by reward history. Using a population-level version of the circular trial permutation test, we found that whole-trial population activity is positively modulated by reward history (*p* = 0.0008, Figure 7A) at a level consistent with our predictions (expected increase in firing from a mean reward of 1*/*2 to 1 of *∼*0.05 Hz neuron^*−*1^, see “Serotonin slowly learns value” above, compared with 0.11 Hz neuron^*−*1^ in our synthetic population based on fitted 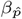 = 1.64 spikes neuron^*−*1^ trial^*−*1^). Encoding of reward history was nearly linear during both the pre-trial baseline and cue epochs (Figure 7B, weighted *r*^2^ = 0.802 during baseline and *r*^2^ = 0.656 during cue), suggesting that tonic and phasic firing participate equally in value prediction. To determine whether a different model could better explain population coding of reward history, we used repeated five-fold cross validation to assess how well mean reward, reward variance, and null models could predict serotonin neuron population activity during the baseline and cue epochs of held-out trials (Figure 7C). Consistent with value prediction, the mean reward model exhibited five- to ten-fold better performance than variability-based alternatives (Table 2). Thus, value prediction describes the effects of reward history on serotonin neuron population activity with a precision considerably better than 0.1 Hz neuron^*−*1^.

**Table 2.** Performance statistics for models of population activity modulation by reward history. Related to Figure 7C.

| Model | Training $r^2$ | | Validation $r^2$ | |
| --- | --- | --- | --- | --- |
|  | Baseline | Cue | Baseline | Cue |
| Value pred. (ours) | $0.894 \pm 0.002$ | $0.713 \pm 0.006$ | $0.763 \pm 0.030$ | $0.526 \pm 0.074$ |
| Variance | $0.111 \pm 0.001$ | $0.208 \pm 0.005$ | $0.088 \pm 0.019$ | $0.111 \pm 0.048$ |
| Std. dev. | $0.109 \pm 0.001$ | $0.168 \pm 0.004$ | $0.086 \pm 0.018$ | $0.085 \pm 0.040$ |
| Entropy | $0.111 \pm 0.001$ | $0.188 \pm 0.004$ | $0.088 \pm 0.019$ | $0.098 \pm 0.044$ |
| Null | 0.000 | 0.000 | 0.000 | 0.000 |

The population activity dynamics shown in Figure 7A exhibit a cue-associated peak, elevated firing during the trace period, and falling activity during the reward, qualitatively consistent with value prediction. Value (without adaptation), surprise, and reward signals are each missing at least one of these properties (Figure 7D left). As a result, quantitative models based on these ideas (see Methods) poorly explain population activity in comparison with value prediction (Figure 7D middle and right). The only tested model that offered performance competitive with value prediction was a surprise signal with added adaptation (mean validation weighted *r*^2^ = 0.749 *±* 0.007 for value prediction and *r*^2^ = 0.642 *±* 0.023 for surprise with adaptation; mean *±* SD of ten cross-validation repeats). However, since the fitted surprise signal with adaptation does not exhibit the expected decrease in activity during the trial (see Methods), the justification for adding adaptation to a model that is already an idealized form of adaptation is dubious, and surprise does not readily explain the effects of reward history on activity (Figure 6 and Figure 7A–C), nor serotonergic responses to punishments (Figure 4) in addition to offering lower performance than value prediction (Figure 7D), we believe it can be rejected. In sum, the fast activity dynamics of serotonin neurons are better explained by value prediction than surprise, reward, or a raw value signal.

Overall, we conclude that value prediction provides a remarkably precise and complete account of the population activity patterns of serotonin neurons during trace conditioning, as foreshadowed by our results at the level of individual neurons.

## Discussion

The *in vivo* activity patterns of serotonin neurons are notoriously difficult to explain. In this work, we show that a time-dependent estimate of cumulative future reward predictively encoded through spike-frequency adaptation, which we call value prediction (Figure 2), unifies a surprisingly wide range of puzzling observations and conflicting theories from the serotonin literature (Figure 1 and Table 1). In particular, phasic activation by reward-predicting cues and primary rewards is explained by a rapid increase in proximity to reward, activation by punishments is explained by a rapid increase in value as the end of the punishment approaches, and tonic firing that reflects reward and punishment context is consistent with a code for the value of upcoming trials (Figure 3). By simulating trace conditioning experiments with different reward sizes and trial durations, we have shown that the appearance of surprise (13) and salience (5) tuning emerges naturally in our model (Figure 4), providing an intuitive link between tuning features that previously seemed conceptually unrelated. To add weight to these qualitative results, we re-analyzed a dataset of serotonin neuron responses to *in vivo* rewards (23). We observed small modulations in serotonin neuron firing by recent reward history, quantitatively consistent with value estimation over hundreds of trials (Figure 6; see also ref. 13). Finally, we directly compared value prediction against competing theories. We found that our theory provides a remarkably precise description of both trial-to-trial changes in activity associated with reward history and within-trial activity dynamics, usually exhibiting predictive performance several times better than alternative models (Figure 7). It has been said that “serotonin’s many meanings elude simple theories” (35). Our work shows that several of these meanings — reward, punishment, surprise, salience, and uncertainty — can be merged into one: value prediction.

### Hiding in plain sight

Why was a serotonergic code for value not established earlier? The idea that serotonin neurons encode value or something very similar is not new (14, 43), but this perspective has fallen increasingly out of favour in recent years because of evidence that seems to directly contradict a value or reward-based code, first and foremost the fact that many serotonin neurons are activated by punishments (11, 13, 44) as well as model-based analysis that suggests that serotonin neurons do not track reward history on the same timescale as changes in foraging behaviour (15). Here we have shown that not only do punishment responses not rule out a value code, they are actually expected if reward-predicting cues evoke transient firing, consistent with experimental results (ref. Fig. 5C in 11). Our work also shows that value coding by serotonin neurons is more precise and temporally extended than behaviour would suggest, again consistent with previous results (13) and adding to an emerging pattern of neural systems having population codes that are more precise than behaviour and perception (45, 46).

Turning to the literature and finding results that seemed puzzling at the time but are predicted by our theory (the apparent reward-specificity of surprise tuning being a notable example; 13) became a recurring theme during this project, leading us to feel that evidence for value coding by serotonin, much like dopamine (47), has long been hiding in plain sight. We hope that future work will uncover more such examples, and, conversely, temper our confirmation bias by highlighting tuning features that are clearly incompatible with value prediction.

### The meaning of predictively-encoded value

An adapting value signal is not the same as value itself, raising interesting questions about how DRN output might be interpreted by downstream regions (and scientists!).

One possibility, although perhaps not a very exciting one, is that value prediction is simply reversed to recover the original value signal. The exceptionally large axonal arborizations of serotonin neurons (48) likely place significant metabolic constraints on the activity of serotonin neurons, and predictive coding through adaptation could act as a sort of compression scheme (21, 22, 33, 49, 50) allowing the serotonin system to broadcast a signal widely using a minimum number of spikes. If adaptation compresses the value signal, how is later decompressed? In theory, the exact answer is simple: leaky integration (Appendix H). In light of the close relationship to predictive coding seen as a compression scheme, it is interesting to note that many of the biological processes involved in decoding serotonin neuron activity implement leaky integration (*e*.*g*., accumulation of serotonin in the extracellular space, slow kinetics of G-protein coupled receptors, and the membrane voltage dynamics of downstream cells). Adapting value might be more similar to value than it first appears.

A more intriguing possibility arises if adaptation already visible in the spiketrains of serotonin neurons is further enhanced downstream, for example via depressing synapses. In that case, our work implies that DRN output would be decoded as the rate of change of value (10). This quantity is required to calculate real-time RPEs (as is value itself; 51, see also 7), and one of the main predictions of the dopaminergic RPE hypothesis of Schultz *et al*. (1) was that the dopamine system should receive input from some region (or collection of regions) that encodes the rate of change of value. Since the dopamine system is one of the main targets of the DRN (52), might serotonin play an important part in computing RPE?

### Behaviour

If serotonin predictively encodes value, how is this signal used to drive learning and behaviour? The ways in which value is used in RL are varied and the effects of serotonergic manipulations are perplexing, but here we offer some speculation.

One of the better established roles of serotonin in regulating behaviour is that fast optogenetic activation of serotonin neurons promotes maintenance of behaviours directed at obtaining imminent rewards, an effect commonly called patience or persistence (16, 17, 53, 54). If serotonin encodes an estimate of relatively immediate reward (due to heavy discounting) that is compared against a longer-term average reward rate, then control policies based on optimal giving up (55) or option interruption (56) could produce similar behaviour. In principle, persistence could also be explained by immediate reinforcement of an ongoing reward-seeking action through policy gradient learning (26) or boosting RPEs (see previous section), but these ideas are difficult to reconcile with evidence that stimulation of serotonin neurons is not generally reinforcing (18, 53, 54, 57, but see 58).

Selfridge’s run and twiddle model (32) offers a more elegant potential explanation of persistence in terms of value. According to this simple control model, the current action is maintained as long as value is increasing (“run”), offering an interesting connection to the predictive coding component of our theory, whereas a decrease in value triggers a random action (“twiddle”). Given the evolutionarily-ancient origins of the serotonin system (59) and its involvement in regulating very coarse aspects of behaviour such as the tradeoff between exploration (taking random or sub-optimal actions) and exploitation (taking actions that are expected to lead to reward) (60), we are tempted to speculate that a relatively primitive control policy might provide the best account of the role of serotonin in behaviour. From that perspective, run and twiddle, which was originally conceived to explain the foraging behaviour of bacteria, might be a good place to start.

### Learning

Confusingly, activation and inhibition of serotonin neurons both promote maintenance of reward-seeking behaviour. However, whereas optogenetic activation of serotonin neurons produces behaviour directed at obtaining immediate rewards that is usually framed in a positive light, chemogenetic inhibition reduces the rate of abandonment of depleted sources of reward (13, 15), which is framed as perseveration. This effect has been explained in terms of a selective decrease in the rate of learning from reward omissions (13, 15). There is a normative reason for the learning rate to decrease when rewards are intrinsically variable (Appendix C) and increase when the environment is non-stationary, prompting the development of models that separately track variance and volatility to enhance learning (61, 62). Serotonin has been proposed to modulate the rate of learning from reward omissions via surprise or uncertainty (13, 15), but, as we have shown, these results are more consistent with value. Finally, a model-based analysis showed that optogenetic manipulations of serotonin neuron activity affected behaviour in a way that was consistent with an enhancement of learning rate (18), but this effect was specific to long timescales and also explained relatively well by RPE boosting (ref. Fig. S14; see previous section). Since all components of RL models affect the rate of change of behaviour, it is plausible that many of the apparent qualitative effects of serotonin on learning could be explained through the lens of value.

### Mechanistic basis of value prediction

The question of where value prediction comes from can be broken into two parts: where does the value signal input originate, and where does predictive coding through adaptation occur?

Since the DRN receives input from nearly the entire forebrain (63), it is unlikely that a single upstream region is completely responsible for computing value, simply because this input would be drowned out by unrelated information streaming into the DRN. Instead, we believe it is likely that the net value input required by our theory is assembled from multiple sources, for example temporally-extended action values from the mPFC (64), and various aspects of reward, including inverted RPE, from the lateral habenula and lateral hypothalamus (45, 65, 66). These possibilities are merely speculation. Our theory does not depend on whether the net value input originates in one particular region or is distributed across many others.

As for the predictive coding aspect of our theory, the exceptionally strong spike frequency adaptation of serotonin neurons is not only sufficient (10), it was the main motivation for the present work. This adaptation comes from a combination of apamin-sensitive potassium currents and spike-triggered changes in spike threshold in individual serotonin neurons (10, 67) as well as network-level recurrent inhibition via 5-HT_1A_ receptors (67, 68). Feed-forward inhibition (63, 69) does not seem to be functionally involved (10). Setting aside this physiological evidence, value prediction on its own technically does not exclude the possibility that at least some of the adaptation visible in the spiketrains of serotonin neurons originates upstream of the DRN.

Selectively inhibiting certain DRN inputs or pharmacologically reducing adaptation *in vivo* could shed light on the mechanisms of value prediction in the serotonin system.

### Phasic and tonic firing

The idea that serotonin neurons encode essentially unrelated signals in phasic and tonic components of their firing rates is popular in the serotonin literature (3, 7–9). Here we have shown that the responses of serotonin neurons to rewards and punishments are well-described by a simple model that does not distinguish between different types of firing. Instead, our model shows that firing patterns that have traditionally been called phasic can be interpreted as increases in firing rate that are short-lived due to adaptation. While we cannot rule out the possibility that serotonin neurons multiplex different quantities in their firing rates in other tasks, it is important to note that the phasic/tonic separation has historically been partly rooted in speculation (7, 8) and the difficulty of formulating a consistent interpretation of serotonergic responses to rewards and punishments (11). Functionally distinct phasic and tonic firing is an exciting hypothesis, but perhaps no longer a good default modelling assumption.

### Heterogeneity

Serotonin neurons are biochemically, developmentally, anatomically, and to some extent electrophysiologically heterogeneous (6). They are probably computationally heterogeneous as well, but in what sense? In principle, serotonin neurons might be quantitatively computationally heterogeneous, meaning that differences in their activity patterns can be captured by adjusting the parameters of our value prediction model, or qualitatively computationally heterogeneous, meaning that value prediction simply does not apply to all serotonin neurons. Our results cannot differentiate between these two possibilities (which are not mutually-exclusive in any case). While value prediction dominates the population activity patterns of serotonin neurons in the data we re-analyzed, many individual neurons seemed essentially unresponsive to the task. If serotonin neurons encode value subject to a wide range of discounting timescales (quantitative heterogeneity), echoing distributional coding in the dopamine system (47, 70, 71), the cells that appear unresponsive might simply exhibit discounting and/or learning rates that are very slow relative to the structure of the experiment (see Figure 8). Consistent with the idea that this experiment is too fast to engage value prediction in most serotonin neurons, the neurons with the clearest responses to reward-predicting cues exhibited reward history modulations much larger than expected based on previous work (see effect size calculation in results). At the same time, the null responses we observed could just as easily be explained by the idea that the neurons in question do not predictively encode value (qualitative heterogeneity). Value prediction subsumes several previous ideas about serotonergic function and is sufficient to explain a surprisingly wide range of results reported in literature, but this does not imply that it is universal.

In view of the marked heterogeneity of the serotonin system in nearly every aspect of its biology, it seems likely that computational heterogeneity in this system is at least partly qualitative. Indeed, there are differences in the tuning features of serotonin neurons across DRN subregions that are difficult for our model to explain (4), as are reported activation by punishment-predicting cues and preference for smaller rewards (11). Perhaps these observations are the result of competing ON and OFF value prediction pathways in the DRN (67) or something else entirely. By testing value prediction against a battery of alternative models, our work provides a template for assessing computational heterogeneity in the serotonin system.

### Top-down meets bottom-up

Here we build on a theme of some of our previous work by adding biological details to a simple model in order to improve its interpretability and performance (10, 72, 73). Here, the success of our model hinges on combining the normative idea of a value signal with spike-frequency adaptation from a bottom-up model of the DRN (10). Neither of these are new to the serotonin field (14, 43, 44, 74–80), but, to the best of our knowledge, they have not previously been combined. The fact that serotonergic responses to rewards and punishments only become interpretable after accounting for the effects of adaptation illustrates the usefulness of elements of biological detail even in normatively-focused branches of computational neuroscience.

### Conclusion

Here we present a simple theory for serotonin’s many meanings: reward, surprise, salience, and uncertainty are different faces of a predictive code for value. Our work links the biology of the serotonin system to a normative account of serotonergic responses to rewards and punishments through value, a quantity that is central to RL theory. On an intuitive level, our definition of value as the expectation of future reward is akin to optimism, providing a conceptual link to the use of serotonergic medications in treating mood disorders. With value prediction, we establish serotonin as “a neural substrate of prediction and reward” (1).

## Methods

### True value signal

We define the true value of a state *s* to be the expected total future reward to be collected by an agent that begins in that state at time *t* and transitions to future states *S*_*t*+1_, *S*_*t*+2_, … according to the dynamics of the environment. Future rewards are discounted by a factor 0 *≤ γ ≤* 1 such that rewards after *t* + 1 are ignored if *γ* = 0 and all rewards are equally valuable if *γ* = 1. This textbook definition of value can be written as an explicit sum over future rewards

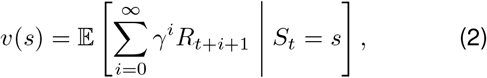

or as an equivalent Bellman recursion

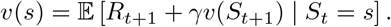

For a trace conditioning experiment with exponentially-distributed inter-trial interval (ITI) durations of mean *L*_ITI_, fixed cue, delay, and reward durations *L*_cue_, *L*_delay_, *L*_reward_, and fixed reward size, the normalized continuous-time true value signal is given by

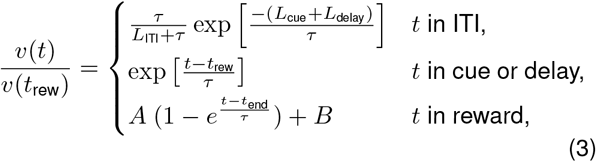

where *τ* = *−dt* ln *γ* is the discounting timescale, *t*_rew_ = *t*_0_ + *L*_cue_ + *L*_delay_ is the start of the reward epoch (given the start of the trial *t*_0_), *t*_end_ = *t*_0_ +*L*_cue_ +*L*_delay_ +*L*_rew_ is the end of the reward epoch, and *A, B* are scaling and offset factors to ensure continuity. (See Appendix D for derivation.) By construction, this normalized definition of the value signal is proportional to the true value signal for any reward size (including negative rewards corresponding to punishments) and can therefore be multiplied by a constant to accommodate different mean rewards.

Note that this model has only one free parameter: the discounting timescale.

### Estimated value signal

To simulate the estimated value signal 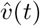 in a trace conditiong experiment, we used the true online TD(*λ*) algorithm of van Seijen *et al*. (40, 81), which is designed to agree more closely with the forward view of TD learning (Figure 2D) than other TD(*λ*) algorithms. van Seijen’s algorithm applies to causal linear value function approximation 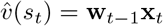, where **x**_*t*_ is a vector of state features and **w**_*t−*1_ are weights from the previous time step (since **w**_*t*_ depends on quantities in the future).

Weights are learned online according to

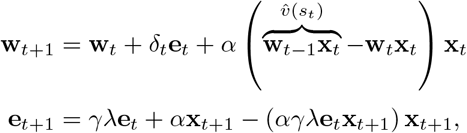

where **e** is the eligibility trace vector and 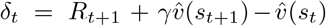 is the reward prediction error (RPE). Compared with traditional TD(*λ*) with eligibility traces, this algorithm adds a correction to the weight update and uses Dutch eligibility traces, which are intermediate to accumulating (**e**_*t*+1_ = *γλ***e**_*t*_ + *α***x**_*t*_) and replacing traces (**e**_*t*+1_ = *γλ***e**_*t*_ ⊙ (1 *−* **x**_*t*_) + *α***x**_*t*_, where *x*_*t*_ is an indicator vector). Following the notation of van Seijen et al. (81), we place the learning rate *α* inside the eligibility trace update rather than in front of *δ*_*t*_ in the weight update.

Simulation was performed using tabular features **x**_*t*_ = **1**_*s*_, eligibility trace *λ* = 0.995, discounting factor *γ* = 0.99, learning rate *α* = 0.01, and time step *dt* = 50 ms.

### Value prediction model

The DRN rate model-based value prediction model is defined as

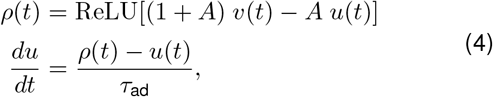

where *ρ*(*t*) is the firing rate of DRN serotonin neurons; *v*(*t*) is the time-dependent net input, assumed to be a value signal (either true or estimated, as indicated in main text); *u*(*t*) is adaptation; *A* and *τ*_ad_ are the strength and timescale of adaptation, respectively; and ReLU[*x*] = max(*x*, 0) is the rectified linear function. The input is rescaled by a factor 1 + *A* so that *ρ*(*t*) = *C* for any constant input *x*(*t*) = *C* independent of the strength of adaptation *A*.

We used *A* = 3 and *τ*_ad_ = 1 s to achieve effective adaptation amplitude and kinetics similar to those observed in our previous experimentally-constrained semi-biophysical model (10). Note that the effective adaptation kinetics are faster than *τ*_ad_ due to feedback between *u*(*t*) and *ρ*(*t*). Adaptation dynamics were numerically integrated using the second-order Runge-Kutta method. Surprise/salience tuning simulations were carried out using a time step *dt* = 1 ms, all others used *dt* = 50 ms.

For a true value signal, the value prediction model has only three free parameters: the discounting timescale, the strength of adaptation, and the adaptation timescale.

### *In vivo* experiment

The dynamic trace conditioning experiment analyzed here has been reported previously by Grossman *et al*. (15). A brief summary is as follows:

Tetrode recordings of optogenetically-tagged serotonin (SERT-expressing) neurons were collected from head-fixed and water-restricted C57BL/6J mice presented with odour-cued water rewards. Each trial consisted of a 1 s odour cue followed by a 1 s delay and a 3 s window during which a fixed-size water reward (approx. 2 µL to 4 µL) could be collected from a lick spout. After the 3 s reward interval, the lick spout was retracted and any remaining water removed via vacuum for 1 s. Inter-trial interval durations were exponentially-distributed with a mean parameter of 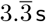 (actual: 3.31 *±* 3.46 s, mean *±* SD, range 0 s to 54.28 s).

Rewards were delivered probabilistically according to a hierarchical Bernoulli point process

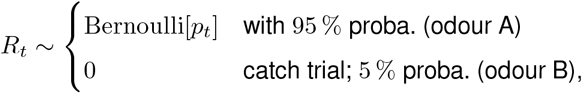

where the reward probability *p*_*t*_ varied according to a block structure. Block lengths were uniformly distributed between 20 and 70 trials. Depending on the recording session, reward probabilities were set to *p*_*t*_ *∈ {*0.2, 0.5, 0.8*}* or *p*_*t*_ *∈ {*0.2, 0.8*}*. Catch trials and trials in which a reward was available but not collected were deemed unrewarded for the purposes of our analysis.

Mice collected 0.45 *±* 0.04 rewards per trial (mean *±* SD; range 0.37 to 0.54) and completed 346 *±* 88 trials per session (range 180 to 565). Mice failed to collect available rewards 4.1 *±* 4.3 % of the time (median 3.4 %, range 0.0 % to 19.0 %). Sessions lasted approximately 1 h (time from start of first trial to start of last trial: 53.1 *±* 12.9 min, range 29.0 min to 88.1 min). Sessions with more trials were not significantly associated with higher or lower reward rates (Pearson *r* = 0.206, *p* = 0.293, *N* = 28 sessions). Data were gathered from five mice (four male, one female) across 28 sessions with 1.32 *±* 0.54 neurons recorded per session (range 1 to 3).

Surgical and experimental procedures were approved by the Johns Hopkins University Animal Care and Use Committee and performed in compliance with the *National Institutes of Health Guide for the Care and Use of Laboratory Animals*.

### Data analysis

#### Reward history

Reward history was operationally defined as the mean reward across the past five trials

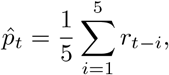

which is an unbiased estimate of the true time-varying Bernoulli reward probability. To mitigate boundary effects, we set *r*_*t*_ = 0.45 for *t <* 0.

Note that the mean reward can be used as a crude proxy for value because the true value is proportional to the true reward probability. The connection between mean reward and true value is also exploited by temporal difference learning methods, which implicitly define value as an exponential moving average of past rewards (Appendix B). We choose to use the five-trial mean reward instead of the true reward probability or a TD estimate so that the connection between the activity of serotonin neurons and the animal’s estimate of a reward statistic is clear.

#### Reward variability

We quantified reward variability using three statistics that can be derived from the estimated Bernoulli reward probability 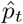: reward variance

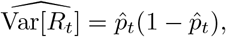

standard deviation

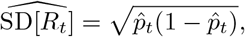

and entropy

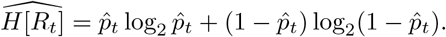

Variance is the focus of our analysis because it is proportional to the absolute RPE used as a measure of surprise or uncertainty in some reward learning models (Appendix F). The other statistics are nearly proportional to variance and are included only to illustrate that our results to not depend on technical details of the variability measure.

#### Quantification of activity

We quantified neural activity using either the peri-stimulus time histogram (PSTH) with 500 ms window width or, as a more coarse-grained metric, the mean number of spikes in a 7.5 s window around each trial (1.5 s pre-trial baseline, 5 s trial, 1 s post-trial baseline), which we refer to as “whole-trial activity”. Baseline activity was defined as the mean of the PSTH 1 s before the start of each trial or, similarly, the number of spikes in a 1 s period just before the start of the trial. Cue-associated activity was defined as the extremum (usually maximum) of the PSTH during the 1 s cue period. PSTHs were not smoothed.

All activity metrics were precision-weighted across neurons and reward history conditions as applicable. For example, the population PSTH for the reward probability 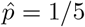 condition was calculated by summing spiketrains from all trials of all neurons where 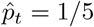 and dividing by the total number of trials being summed. The resulting population PSTH can be seen as a weighted average of the PSTHs of individual neurons, where each neuron is weighted according to the precision (inverse variance) of its PSTH *N/σ*^2^.

#### Dynamical model definitions

##### Value

The value prediction and value models of serotonin neuron activity are defined by Eq. **??** and 4. The discounting timescale, adaptation strength, and adaptation timescale are estimated from the data along with scale and offset parameters.

##### Surprise

Surprise is defined in reward learning as the absolute reward prediction error |*δ*_*t*_| and in information theory as log_2_ *p*, both of which are zero for deterministic events and greater than zero for stochastic events. Therefore, each instant in the inter-trial interval, the beginning of the trial, and the beginning of reward delivery all have non-zero surprise, while all other moments during the trial have zero surprise.

Our surprise-like model is defined as a piecewise constant function

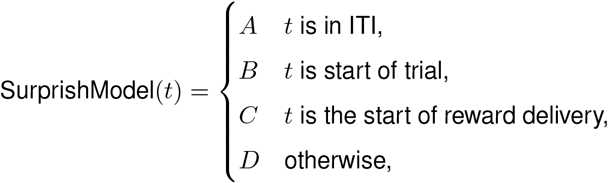

subject to the restrictions

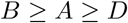

and

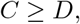

where the coefficients *A, B, C, D* are estimated from the data.

##### Reward

The reward model is a piecewise constant function

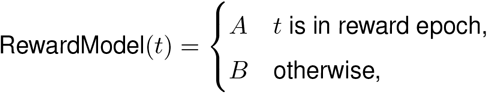

where the coefficients *A, B* are estimated from the data.

##### Null

The null model is a constant function with an offset parameter estimated from the data.

##### Surprise with adaptation

The surprise-like model with adaptation is modified from the surprise-like model defined above:

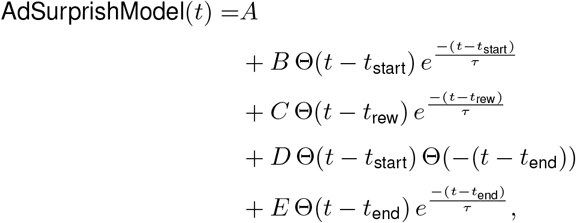

subject to the restrictions

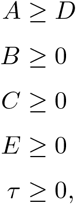

where Θ(*·*) is the Heaviside step function and the coefficients *A, B, C, D, E, τ* are estimated from the data. As before, *A, B, C, D* represent the activity during the ITI, trial start, reward start, and the remainder of the trial, respectively. *E* represents the amplitude of the overshoot at the end of the trial and *τ* is the timescale of adaptation.

While adaptation can be used to compute surprise, we are doubtful that adaptation should be added to a model of surprise itself: the fact that adding adaptation actually slows down the kinetics of the surprise signal is a major conceptual difficulty. We include this model in our analysis only for completeness.

#### Reward with adaptation

The reward model with adaptation is modified from the reward model defined above:

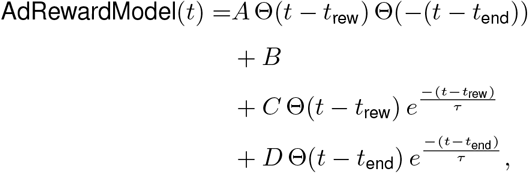

subject to the restrictions

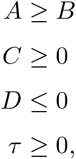

where the coefficients *A, B, C, D, τ* are estimated from the data. As before, *A, B* represent the activity during the reward period and at all other times, respectively. *C* parameterizes the amplitude of the phasic activity associated with reward onset, *D* parameterizes the amplitude of the undershoot associated with reward offset, and *τ* is the time constant of adaptation.

#### Dynamical model fitting

Before fitting, all models were smoothed with a 500 ms boxcar filter to simulate a PSTH and lagged by 150 ms to account for perceptual delays.

All models were fitted by minimizing the mean squared error on the population PSTH (see Quantification of activity). For the reward model (without adaptation) this was accomplished using linear regression. For all other models, this was accomplished using bounded/constrained gradient-based optimization methods provided by `scipy.optimize.minimize` (L-BFGS-B or sequential least-squares quadratic programming).

Performance was assessed using repeated five-fold cross validation. Data was stratified by neuron identity (which partially reflects reward history beyond the five-trial horizon, Figure 6E) and reward history level prior to assigning folds in order to minimize class imbalances between training and validation sets.

We use cross validation rather than AIC or BIC because cross validation does not rely on distributional assumptions that would be needed to formulate a likelihood function for each model.

#### Analysis of reward modulation

Reward modulation was defined as the slope 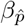 of a regression line between an activity metric (see Quantification of activity) and the estimated reward probability

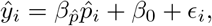

where *y*_*i*_ is the measured activity, 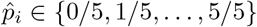 is the estimated reward probability (see Reward history), *β*_0_ is the intercept of the regression line, and *ϵ*_*i*_ is a residual. The regression model was fitted using weighted least-squares (observations were precision-weighted; see Quantification of activity).

Statistical significance of the slope 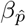 was assessed using circular trial permutation or bootstrapping.

Circular trial permutation tests are not sensitive to autocorrelations in timeseries data that can significantly increase the false positive rates of classical and shuffling-based statistical tests (41). These tests were performed by shifting the per-trial estimated reward probabilities 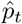 to break the alignment between activity and reward history while controlling for other correlations in the data. For example, the reward probabilities for a *T* trial experiment 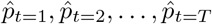 can be shifted *D* places to generate un-aligned probabilities

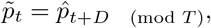

and the analysis described above can be repeated using the shifted probabilities 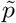. The value of the slope 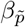 obtained using this procedure represents the apparent reward modulation under the null hypothesis that activity and reward history are not actually related (because the trials were shifted). In the case of whole-trial reward modulation of population activity, the permutation procedure was repeated 2500 times to generate a distribution for 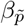 used to obtain an approximate *p*-value for 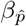. In the case of whole-trial reward modulation of individual neurons, the procedure was repeated exhaustively to generate exact *p*-values. In both cases, we restricted 10 *≤ D ≤ T −*10 due to very high experimental design-related autocorrelations in 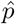 (specifically, block structure and the fact that 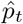 and 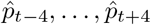 are calculated on overlapping sets of trials). Removing this restriction *post hoc* did not meaningfully affect our results.

Bootstrap distributions for activity metrics *ŷ*_*i*_, regression predictions *y*_*i*_, and regression slopes 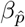 were generated by sampling trials 1000 times with replacement within reward probability levels 0*/*5, 1*/*5, …, 5*/*5 and neurons as applicable. Activity metric distributions were corrected for Monte-Carlo bias (82). Statistical significance of the regression slope 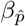 was assessed using the 99 % confidence interval method.

For consistency with dynamical model comparison, reward modulation models based on reward history 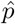 and related statistics Var[*R*], SD[*R*], *H*[*R*] (see corresponding section above) were compared using repeated stratified five-fold cross validation. Performance is presented as the mean variance explained across folds for each repeat.

### Statistical analysis

Statistical tests are specified in the main text and methods above. Non-parametric tests were used as much as possible; where samples sizes were so small that the loss of power associated with non-parametric tests became prohibitive, robust tests were used. All tests are two-sided unless otherwise stated. Sign tests were one-sided because our theory implies that the relevant effects should have a specific sign. Exact *p* values are reported in the main text. Results were considered statistically significant at *p ≤* 0.05 and *p ≤* 0.1 was considered a trend. Sample sizes were not predetermined because only previously published data was used. *N* = 37 neurons in nearly all cases. Because neurons were usually recorded individually (see “*In vivo* experiment” above), we considered them to be independent biological replicates for the purposes of statistical analysis. Error bars represent 95 % confidence intervals. Uncertainties are presented as standard deviation in the main text unless otherwise specified.

### Data and code availability

Previously-published data is available on the Dryad repository (23). Code will be made available on GitHub.

### Copyright permissions

Elements of Figure 3B and C, Figure 4C, and Figure 5 have been reproduced from Cohen *et al*. (11), Matias *et al*. (13), and Zhong *et al*. (12), respectively, under the Creative Commons Attribution license (CC-BY 4.0). A bar chart from Paquelet *et al*. (5), which is covered by the Creative Commons Attribution Non-commercial No Derivatives license (CC-BY-NC-ND 4.0), has been included in Figure 4D with kind permission from Bradley Miller.

Artwork has been modified by changing colours, replacing axis annotations with icons or larger text, and/or adding schematics to improve clarity. Due to space constraints, small reward and neutral stimulus groups were removed from the vignette in Figure 4C, as were statistical annotations. In all cases, our aesthetic modifications do not change the interpretation of the underlying data. References to the specific figures from the original publications are included in our captions.

## Acknowledgements

We wish to acknowledge that this work was carried out on the unceded and unsurrendered land of the Algonquin Anishinaabe people. The data we re-analyzed was collected on the unceded lands of the Piscataway and Susquehannock people.

Emerson Harkin is grateful for PGS-D and Queen Elizabeth II Scholarships in Science and Technology awards from the Natural Sciences and Engineering Research Council and Government of Ontario, respectively. This work was funded by grants to Richard Naud.

We thank Bradley Miller and Paul Albert for providing feedback on an initial draft of this paper. We also thank Michael Lynn and Sébastien Maillé for many helpful brainstorming sessions and input on figure design; John Beninger for assisting with troubleshooting population-level analysis; and all members of the Béïque and Naud labs for helpful discussions.

## Author contributions

Emerson Harkin conceptualized the project, created the model, performed all simulations and data analysis, performed mathematical analysis, created all figures, and wrote the first and final drafts of the manuscript as well as all drafts of the discussion section. Richard Naud provided supervision, funding, and extensive input on all aspects of the project as well as performing mathematical analysis and writing the intermediate draft. Jean-Claude Béïque provided extensive input and helpful discussion throughout the project, as well as detailed comments on the manuscript. Cooper Grossman and Jeremiah Cohen provided data and validated the design of our analysis. Cooper Grossman provided helpful discussion and extensive input on comparisons with the uncertainty model, significantly strengthening this aspect of the work. Jeremiah Cohen provided helpful discussion and detailed input on the manuscript.

## A Connection between state value *v*(*s*) and state-action value *q*(*s, a*)

It is well known that the state-action value depends on the state value: *q*(*s, a*) = 𝔼 [*R*_*t*+1_ + *γv*(*S*_*t*+1_) | *S*_*t*_ = *s, A*_*t*_ = *a*] (25). As we show below, the state value can also be expressed in terms of state-action values.

Following the notational conventions of RL theory, let *S, A* be random variables denoting the state and action and let *s, a* be a specific state and action. Let *v*(*s*) and *q*(*s, a*) be the state and state-action value functions, respectively, under the action-generating policy *π*(*a* | *s*) = Pr[*A*_*t*_ = *a* | *S*_*t*_ = *s*]. The state value function *v*(*s*) is equivalent to the expected state action value *q*(*s, a*) under the policy *π*:

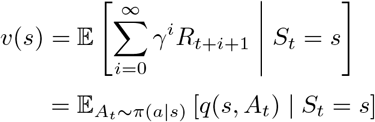

### Proof

The state-action value function is the expected cumulative discounted future reward to be obtained after taking action *a* in state *s*, written as

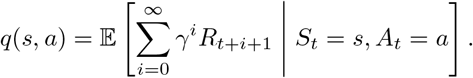

Note that this is exactly the same as the state value *v*(*s*) except for the dependence on the chosen action *A*_*t*_ = *a*.

To prove that *v*(*s*) is the expected *q* value, we need to incorporate *A*_*t*_ into *v*(*s*). To simplify notation, let *G*_*t*_ be the random variable 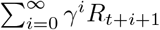. Using *G*_*t*_ and expanding expectation, we can rewrite the state value as follows

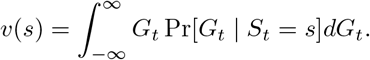

Incorporating the chosen action *A*_*t*_ by conditioning, we obtain

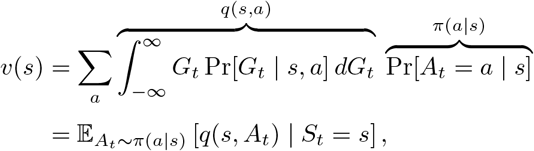

completing the proof.

## B Temporal difference learning averages past rewards

Consider an experiment with a single state (*e*.*g*., a trace conditioning experiment where each trial is considered to be a time step) and a stochastic reward *R ∼* Bernoulli[*p*]. Learning the state value 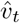 using the RPE 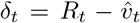 causes the state value to be an exponential moving average of past rewards that is an unbiased estimate of the reward probability *p*

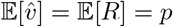

### Proof

The current state value 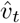 depends on the past state value 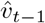 and RPE *δ*_*t−*1_

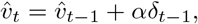

where 0 *≤ α ≤* 1 is the learning rate. Expanding the above, we obtain

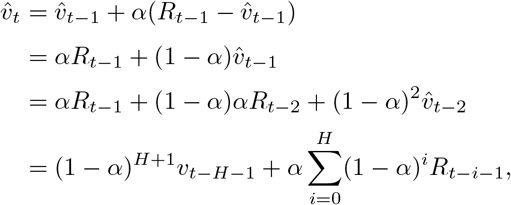

where *H ∈* ℕ is the time horizon. Taking an infinite horizon, the term 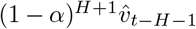 vanishes, and we are left with value as a scaled exponential moving average of past rewards

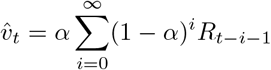

with an estimation timescale *τ*_est_ = *−dt* ln(1 *− α*).

Taking the expectation shows that the above is an unbiased estimate of the reward probability

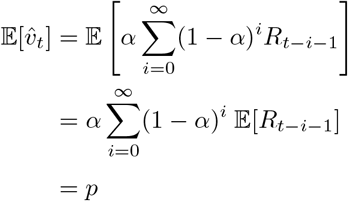

by linearity of expectation 𝔼 [*αX*] = *α* 𝔼 [*X*], simplification of the geometric series 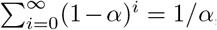, and expectation of the Bernoulli reward 𝔼 [*R*_*t*_] = *p*.

The reader may verify that the above implies that 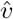 is an *unbiased* estimator by adding a bias term *b* and showing that it is zero: 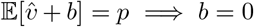. This completes the proof.

## C Learning rate controls value variance

Appendix B shows that the state value 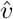 is an unbiased estimate of the mean reward 𝔼 [*R*] = *p*. The variance of this estimate is set by the TD learning rate *α* according to

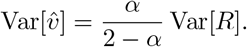

### Proof

ecall that the value estimate can be written as

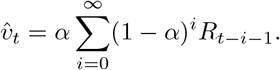

Taking the variance and moving the constant terms out, we obtain

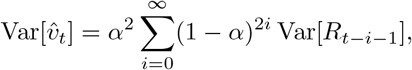

remembering that Var[*α* Σ _*i*_ *X*_*i*_] = *α*^2^ Σ _*i*_ Var[*X*_*i*_] for constant *α* and independent *X*_*i*_. Removing time dependence by assuming Var[*R*_*t−i−*1_] = Var[*R*] *∀t, i* and simplifying the constant term using convergence of the geometric series completes the proof.

## D Derivation of true value in trace conditioning experiments

The normalized true value signal *v*(*t*)*/v*(*t*_rew_) in a trace conditioning experiment is given by the piecewise function

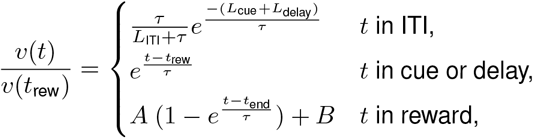

presented in Equation (3). (See Methods for the meaning of each variable.)

To derive this equation, we model a trace conditioning experiment as a Markov reward process (MRP). The states in the MRP are labelled *s*_0_, *s*_1_, *s*_2_, …, *s*_*M*+*N*_, where *s*_0_ is the ITI, *s*_1_, *s*_2_, …, *s*_*M*_ are the trial states representing the *M* time steps between the start of the cue in *s*_1_ and the end of the delay period in *s*_*M*_, and *s*_*M*+1_, *s*_*M*+2_, …, *s*_*M*+*N*_ are the reward states representing the *N* time steps during the reward period. The transition probabilities are

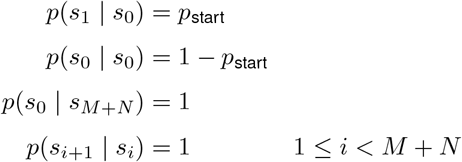

where *p*(*s*_*j*_ | *s*_*i*_) represents the probability of transitioning to *s*_*j*_ at the next time step given that the current state is *s*_*i*_. These transition probabilities were chosen so that the dwell time in the ITI state *s*_0_ follows a geometric distribution with mean 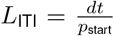, reflecting exponentially distributed ITI durations commonly used in experiments, and so that the fixed durations of the cue, delay, and reward epochs are given by *L*_trial_ = *L*_cue_ + *L*_delay_ = *M dt* and *L*_rew_ = *N dt*. A reward of size *r/N* is delivered in each of the *N* reward states, such that each trial ends in a total reward of size *r*. As noted in the main text, we define the value of each state in terms of the expected discounted future reward

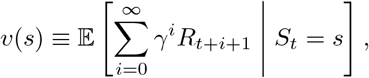

which can also be written in Bellman form as

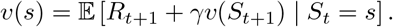

In the three sections below, we show how each part of the continuous time normalized true value signal *v*(*t*)*/v*(*t*_rew_) can be derived from the MRP and value function *v*(*s*) given above in the limit of *dt →* 0.

### True value during the ITI

Writing the value of the ITI state *v*(*s*_0_) using the Bellman form and expanding the expectation shows that it is dependent on itself and the value of the first trial state

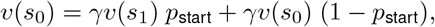

which can be solved in terms of *v*(*s*_0_), yielding

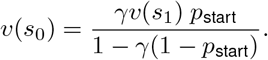

The first trial state *v*(*s*_1_) can be written in terms of the final trial state *v*(*s*_*M*_)

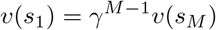

since the state transitions are deterministic and no rewards are delivered during the cue and delay epochs.

Substituting the value of the first trial state *v*(*s*_1_) into the value of the ITI state *v*(*s*_0_) and normalizing by the peak value during the trial *v*(*s*_*M*_) yields

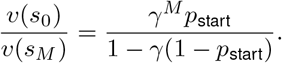

Using *M* = (*L*_cue_ + *L*_delay_)*/dt* and *p*_start_ = *dt/L*_ITI_ from the problem definition and letting 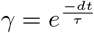, the relative ITI value can be rewritten

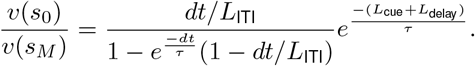

Taking the limit as *dt →* 0 completes the derivation.

### True value during trace and delay epochs

Using the Bellman form of the value function and the fact that no rewards are delivered during the cue and trace periods, the value leading up to reward can be written

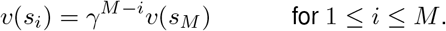

Normalizing by *v*(*s*_*M*_) and converting to continuous time using 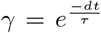 and *M* = *L*_trial_*/dt* completes the derivation.

### True value during the reward epoch

The true value starting in state *s*_*M*_ and continuing through the reward epoch is

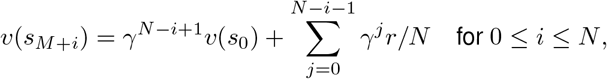

abusing notation by allowing 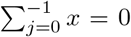. Removing the contribution of *v*(*s*_0_) and normalizing out the reward, we are left with

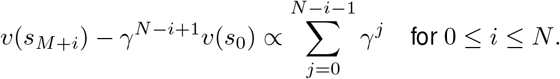

The sum of discounting factors on the right hand side can be rewritten in continuous time as the integral of an exponential discounting kernel

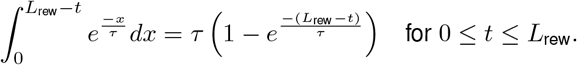

Incorporating the scaling and offset terms *A, B* to ensure continuity with the normalized value function at *v*(*t*_rew_) and *v*(*t*_end_) completes the derivation.

## E Inter-trial interval value reflects value at the start of the next trial

Let the state *a* be that the animal is in the ITI, let the state *b* be that the animal is at the very start of a trial, and let *T* be a random variable that represents the number of time steps remaining in the ITI before the start of the next trial. The true value of the ITI state *v*(*a*) is proportional to the value at the start of the next trial *v*(*b*) as follows

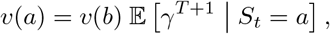

where *γ* is the discounting factor.

### Proof

Recall that the true value of a state *S, s* is defined as the expected sum of exponentially-discounted future rewards

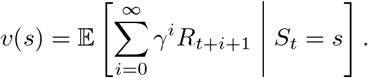

Applying this definition to the states *a, b*, we obtain 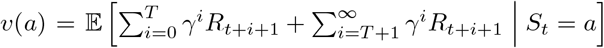 and 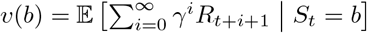, where the sum in *v*(*a*) is split between the value of the rest of the ITI and the discounted value at the start of the next trial. Since the value at the start of the next trial is *v*(*b*), we can rewrite the ITI value as follows

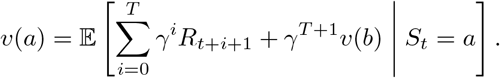

Observing that the cumulative reward during the ITI is zero by definition 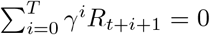, and also observing that *v*(*b*) is a constant that can be factored out of the expectation, we find that the ITI value is

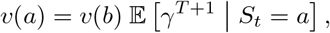

completing the proof.

### Notes

- This proof can easily be extended to the value of any state in the upcoming trial (not just the first state) by redefining *T* and *b*.
- If the ITI durations are drawn from a geometric (or exponential) distribution, then the scaling factor 𝔼[*γ*^*T* +1^ |*S*_*t*_ = *a*] does not depend on the amount of time spent in the ITI so far. The value during the ITI *v*(*a*) is therefore constant.

## F Relationship between uncertainty and reward variance

Absolute RPEs |*δ*| are sometimes used in RL as a measure of variability or uncertainty that can be used to regulate learning (15), but the precise statistical interpretation of |*δ*| is unclear. Here we show that for vanilla TD learning from Bernoulli rewards, the average absolute RPE is proportional to the reward variance

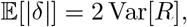

but this relationship can be distorted by asymmetric learning rates and forgetting.

### Proof for vanilla TD

For simplicity, let *R ∼* Bernoulli[*p*], let the RPE be 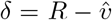, and set value to its fixed point 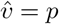 (since 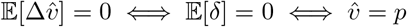 under TD learning). Substituting *p* into the RPE and taking the expectation of the absolute value, we get

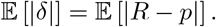

Since the reward is binary *R ∈ {*0, 1*}*, we can easily rewrite the expectation above and simplify

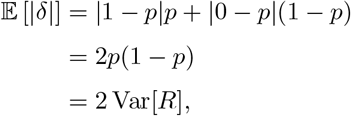

completing the proof.

The relationship between absolute RPE |*δ*| and reward variance shown analytically above holds well in practice even if the value estimate 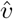 is not generally exactly equal to its fixed point *p* (Figure 13).

**Fig. 13.**
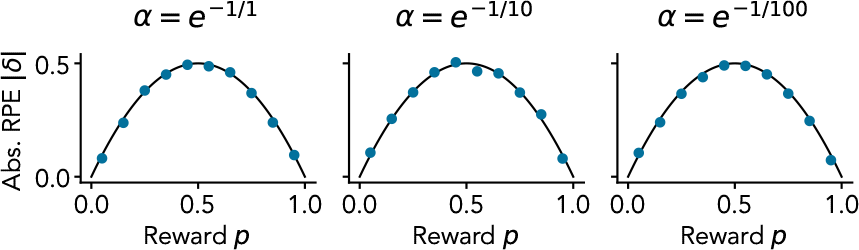
Numerical verification of relationship between absolute RPE |*δ*| and reward variance Var[*R*] under TD learning. Panels show |*δ*| mean *±* SEM (blue points; error bars are too small to be visible) calculated from simulated TD learning trials against 2 Var[*R*] (black lines). Simulations were burned in for 500 trials and RPE statistics were calculated on 2000 trials. Learning rate *α* used in the TD model is indicated at top of each panel.

### Effect of asymmetric learning

In vanilla TD learning, the value estimate is updated proportional to the RPE 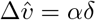, but a common extension is to use different learning rates for positive and negative RPEs

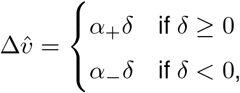

called asymmetric learning. Under asymmetric learning, value has a fixed point that is different from the reward probability 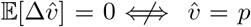. We can find the fixed point by solving

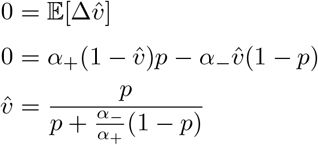

Substituting the above into the expectation of the absolute RPE 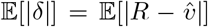, we find that there is no longer a clear connection to the variance of the Bernoulli reward Var[*R*] (Figure 14).

**Fig. 14.**
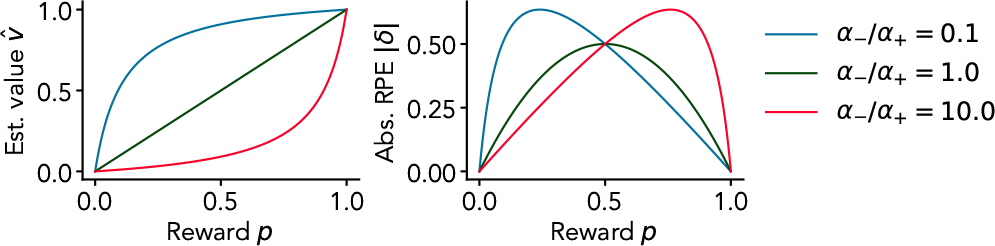
Effect of asymmetric learning rates on value 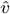 and absolute RPE |*δ*| in TD learning.

### Effect of forgetting

Another common modification of TD learning is to include a forgetting rate 0 *≤ ζ ≤* 1 in the RPE 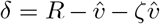. Solving 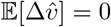 as above, we find the fixed point

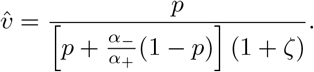

The main effect of forgetting is to decrease the estimated value when the reward probability is high, increasing the absolute RPE in this range (Figure 15).

**Fig. 15.**
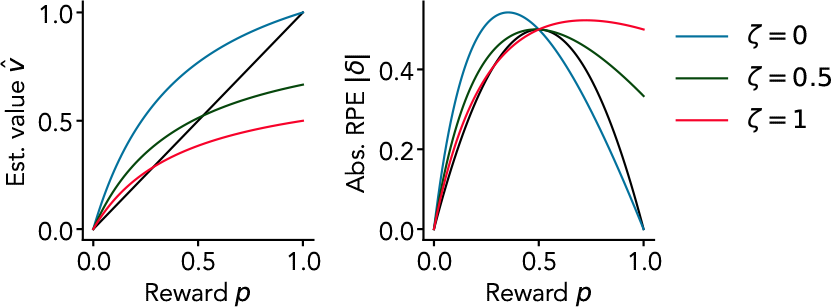
Effect of forgetting on value 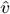 and absolute RPE |*δ*| under asymmetric TD learning with 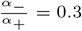.

## G Perturbation analysis

### Setup

According to the value prediction theory, the trial-aligned activity patterns of serotonin neurons in reward learning experiments should resemble the normalized true value signal derived in Appendix D scaled by the reward probability. Therefore, if an animal’s estimate of the reward probability is dynamically changing, then the activity of serotonin neurons should scale up and down accordingly. Assuming that the animal estimates the reward probability on the basis of the proportion of recent trials that were rewarded, the value prediction theory makes a simple testable prediction: in a given experiment, the firing rates of serotonin neurons should be positively correlated to the proportion of recent trials that were rewarded. We do no know, however, the precise timescale over which the recent rewards affect the value function. To circumvent this problem, we use perturbation theory to derive a simpler expression relating the value with the fraction of recent trials that were rewarded.

Our value prediction theory states the firing rate of serotonergic neurons is proportional to the predictively coded value signal. Over the slow timescale of the whole-trial, the predictively coded value signal becomes the value signal.

Our goal is thus to relate the value signal

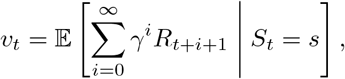

with fluctuations in the recent reward history 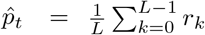, for some small number of recent trials *L* and where *r*_*k*_ refers to the *k*th reward in the past with respect to time *t*.

We begin by proving that 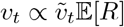, where the normalized value signal 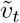 from Appendix D captures within-trial value dynamics and the expected reward 𝔼 [*R*] is responsible for trial-to-trial fluctuations. By definition,

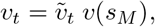

where *v*(*s*_*M*_) is the true value just before reward delivery (see Appendix D), so it remains only to be shown that *v*(*s*_*M*_) *∝* 𝔼 [*R*]. Assuming the reward lasts only one time step, we can write

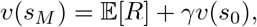

where *v*(*s*_0_) is the true value during the ITI. Using the fact that *v*(*s*_0_) *∝ v*(*s*_*M*_) from Appendix E, we can introduce a temporary proportionality constant *C* and simplify

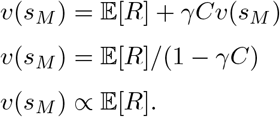

Thus we have that

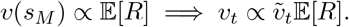

The proof can be extended to rewards that last more than one timestep without much difficulty.

There are multiple ways of calculating the expected reward 𝔼 [*R*], but in TD learning methods, the expected reward is an exponential moving average of past rewards (Appendix B). Therefore, the value signal fluctuates according to an unknown value estimation timescale *τ*_*e*_

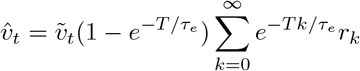

Using the fact that the total trial and ITI duration *T* times the trial horizon *L* is much smaller than the estimation timescale *τ*_*e*_, by first order Taylor expansion of the exponential terms we get

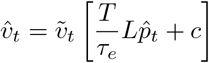

where 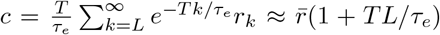 where 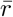 is the average reward on a long time horizon. The relative change in value is then

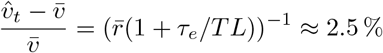

using an estimation time scale of *τ*_*e*_ = 200 trials (12, 13), an *L* = 5 trial horizon, and an average reward rate of 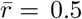. This implies that if the average firing rate of a serotonin neuron is 2 Hz, we expect to observe activity modulations of roughly 0.05 Hz in either direction around the mean based on a five-trial reward history.

### Note

As a rough guide to the sensitivity of this calculation, consider that if the estimation time scale were an order of magnitude faster than what has previously been reported (12, 13) (*τ*_*e*_ = 20 trials), then we would expect activity modulations of roughly 0.8 Hz (40 % of a 2 Hz baseline). Therefore, changes in serotonergic activity associated with a five-trial reward history should be much smaller than the multi-Hz within-trial fluctuations in activity explained by 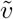 and observed experimentally (11), even if our estimate of the value estimation timescale *τ*_*e*_ is badly wrong.

## H Predictive coding exactly cancels leaky decoding

Let *I*(*t*) be the net input to the DRN, let *f* (*·*) be the encoding function of the DRN, and let *g*(*·*) be the decoding function of some downstream region. The output of the DRN is (*f ° I*)(*t*) and the decoded signal is (*g ° f ° I*)(*t*). Assuming that *g* performs leaky integration on its input, the decoded signal is given as follows

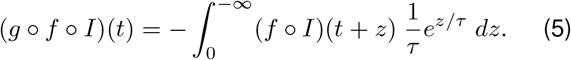

Following our previous work (10), assuming *f* predictively encodes its input such that

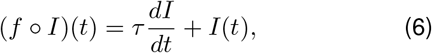

then

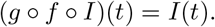

### Proof

Substituting Eq. 6 in Eq. 5 and expanding the resulting integral yields

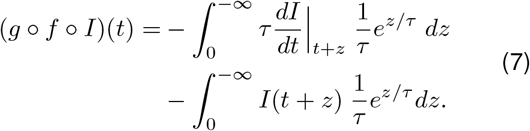

The time derivative of the input 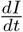 can be removed from the first term using integration by parts, yielding

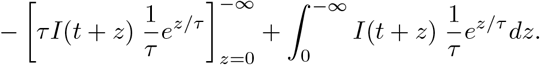

Substituting the above into Eq. 7 causes the remaining integrals to cancel and changes the sign on the non-integral term, giving

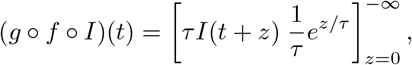

which simplifies to

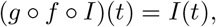

completing the proof.

## Footnotes

1 This does not seem implausible. Trace conditioning trials are typically only a few seconds long−if humans had a discounting timescale on the order of a minute or less, no-one would ever read beyond the first few sentences of this manuscript.

## References

[1] W. Schultz, P. Dayan, and P. R. Montague. A neural substrate of prediction and reward. Science 275, no. 5306, 1593–1599. (1997)

[2] R. S. Sutton and A. G. Barto. Reinforcement Learning, 2nd ed. (The MIT Press, 2018)

[3] C. D. Grossman and J. Y. Cohen. Neuromodulation and neurophysiology on the timescale of learning and decision-making. Annual Review of Neuroscience 45, 317–337. (2022)

[4] J. Ren, D. Friedmann, J. Xiong, C. D. Liu, B. R. Ferguson, T. Weerakkody, K. E. DeLoach, C. Ran, A. Pun, et al. Anatomically defined and functionally distinct dorsal raphe serotonin sub-systems. Cell 175, no. 2, 472–487. (2018)

[5] G. E. Paquelet, K. Carrion, C. O. Lacefield, P. Zhou, R. Hen, and B. R. Miller. Single-cell activity and network properties of dorsal raphe nucleus serotonin neurons during emotionally salient behaviors. Neuron 110, no. 16, 2664–2679. (2022)

[6] B. W. Okaty, K. G. Commons, and S. M. Dymecki. Embracing diversity in the 5-HT neuronal system. Nature Reviews Neuroscience 20, no. 7, 397–424. (2019)

[7] N. D. Daw, S. Kakade, and P. Dayan. Opponent interactions between serotonin and dopamine. Neural Networks 15, no. 4-6, 603–616. (2002)

[8] D. Asher, A. Craig, A. Zaldivar, A. Brewer, and J. Krichmar. A dynamic, embodied paradigm to investigate the role of serotonin in decision-making. Frontiers in Integrative Neuroscience 7. (2013)

[9] K. Wong-Lin, D.-H. Wang, A. A. Moustafa, J. Y. Cohen, and K. Nakamura. Toward a multiscale modeling framework for understanding serotonergic function. The Journal of Psychopharmacology 31, no. 9, 1121–1136. (2017)

[10] E. F. Harkin, M. B. Lynn, A. Payeur, J.-F. Boucher, L. Caya-Bissonnette, D. Cyr, C. Stewart, A. Longtin, R. Naud, et al. Temporal derivative computation in the dorsal raphe network revealed by an experimentally-driven augmented integrate-and-fire modeling framework. eLife 12, e72951. (2023)

[11] J. Y. Cohen, M. W. Amoroso, and N. Uchida. Serotonergic neurons signal reward and punishment on multiple timescales. eLife 4, e06346. (2015)

[12] W. Zhong, Y. Li, Q. Feng, and M. Luo. Learning and stress shape the reward response patterns of serotonin neurons. The Journal of Neuroscience 37, no. 37, 8863–8875. (2017)

[13] S. Matias, E. Lottem, G. P. Dugué, and Z. F. Mainen. Activity patterns of serotonin neurons underlying cognitive flexibility. eLife 6, e20552. (2017)

[14] M. Luo, Y. Li, and W. Zhong. Do dorsal raphe 5-HT neurons encode “beneficialness”? Neurobiology of Learning and Memory 135, 40–49. (2016)

[15] C. D. Grossman, B. A. Bari, and J. Y. Cohen. Serotonin neurons modulate learning rate through uncertainty. Current Biology 32, no. 3, 586–599. (2022)

[16] E. Lottem, D. Banerjee, P. Vertechi, D. Sarra, M. oude Lohuis, and Z. F. Mainen. Activation of serotonin neurons promotes active persistence in a probabilistic foraging task. Nature Communications 9, no. 1, 1000. (2018)

[17] K. Miyazaki, K. W. Miyazaki, A. Yamanaka, T. Tokuda, K. F. Tanaka, and K. Doya. Reward probability and timing uncertainty alter the effect of dorsal raphe serotonin neurons on patience. Nature Communications 9, no. 1, 2048. (2018)

[18] K. Iigaya, M. S. Fonseca, M. Murakami, Z. F. Mainen, and P. Dayan. An effect of serotonergic stimulation on learning rates for rewards apparent after long intertrial intervals. Nature Communications 9, no. 1, 2477. (2018)

[19] K. Doya. Metalearning and neuromodulation. Neural Networks 15, no. 4, 495–506. (2002)

[20] N. Schweighofer, M. Bertin, K. Shishida, Y. Okamoto, S. C. Tanaka, S. Yamawaki, and K. Doya. Low-serotonin levels increase delayed reward discounting in humans. The Journal of Neuroscience 28, no. 17, 4528–4532. (2008)

[21] M. Srinivasan, S. Laughlin, and A. Dubs. Predictive coding: A fresh view of inhibition in the retina. Proceedings of the Royal Society of London 216, no. 1205, 427–459. (1982)

[22] M. Chalk, O. Marre, and G. Tkačik. Toward a unified theory of efficient, predictive, and sparse coding. Proceedings of the National Academy of Sciences 115, no. 1, 186–191. (2018)

[23] J. Cohen, C. Grossman, and B. Bari. Serotonin neurons modulate learning rate through uncertainty. Dryad. (2021) doi: 10.5061/dryad.cz8w9gj4s

[24] R. E. Bellman and S. E. Dreyfus. Applied Dynamic Programming. (Princeton University Press, 1962)

[25] C. Watkins. Learning from Delayed Rewards. (1989)

[26] R. S. Sutton, D. A. McAllester, S. P. Singh, and Y. Mansour. Policy gradient methods for reinforcement learning with function approximation. Advances in Neural Information Processing Systems 12. (1999)

[27] C. Watkins. Modes of Control of Behaviour in Learning from Delayed Rewards, pp. 55–71. (1989)

[28] C. Watkins and P. Dayan. Q-Learning. Machine Learning 8, 279–292. (1992)

[29] R. S. Sutton. Learning to predict by the methods of temporal differences. Machine Learning 3, no. 1, 9–44. (1988)

[30] C. Watkins. Primitive Learning in Learning from Delayed Rewards, pp. 81–113. (1989)

[31] R. J. Williams. Simple statistical gradient-following algorithms for connectionist reinforcement learning. Machine Learning 8, 229–256. (1992)

[32] O. Selfridge. Some themes and primitives in ill-defined systems in Adaptive Control of Ill-Defined Systems, pp. 21–26. (1984)

[33] M. Spratling. A review of predictive coding algorithms. Brain and Cognition 112, 92–97. (2017)

[34] B. N. Lundstrom, M. H. Higgs, W. J. Spain, and A. L. Fairhall. Fractional differentiation by neocortical pyramidal neurons. Nature Neuroscience 11, no. 11, 1335–1342. (2008)

[35] P. Dayan and Q. Huys. Serotonin’s many meanings elude simple theories. eLife 4, e07390. (2015)

[36] K. Miyazaki, K. W. Miyazaki, and K. Doya. Activation of dorsal raphe serotonin neurons underlies waiting for delayed rewards. Journal of Neuroscience 31, no. 2, 469–479. (2011)

[37] J. M. Pearce and G. Hall. A model for pavlovian learning: variations in the effectiveness of conditioned but not of unconditioned stimuli. Psychological Review 87, no. 6, 532–552. (1980)

[38] Z. Liu, J. Zhou, Y. Li, F. Hu, Y. Lu, M. Ma, Q. Feng, J.-e. Zhang, D. Wang, et al. Dorsal raphe neurons signal reward through 5-HT and glutamate. Neuron 81, no. 6, 1360–1374. (2014)

[39] Y. Li, W. Zhong, D. Wang, Q. Feng, Z. Liu, J. Zhou, C. Jia, F. Hu, J. Zeng, et al. Serotonin neurons in the dorsal raphe nucleus encode reward signals. Nature Communications 7, no. 1, 10503. (2016)

[40] H. Van Seijen, A. R. Mahmood, P. M. Pilarski, M. C. Machado, and R. S. Sutton. True online temporal-difference learning. The Journal of Machine Learning Research 17, no. 1, 5057–5096. (2016)

[41] L. Elber-Dorozko and Y. Loewenstein. Striatal action-value neurons reconsidered. eLife 7, e34248. (2018)

[42] L. V. Hedges. Estimation of effect size under nonrandom sampling: The effects of censoring studies yielding statistically insignificant mean differences. Journal of Educational Statistics 9, no. 1, 61–85. (1984)

[43] K.-i. Okada, K. Nakamura, and Y. Kobayashi. A neural correlate of predicted and actual reward-value information in monkey peduncu-lopontine tegmental and dorsal raphe nucleus during saccade tasks. Neural Plasticity 2011, 1–21. (2011)

[44] K. Hayashi, K. Nakao, and K. Nakamura. Appetitive and aversive information coding in the primate dorsal raphé nucleus. The Journal of Neuroscience 35, no. 15, 6195–6208. (2015)

[45] E. L. Sylwestrak, Y. Jo, S. Vesuna, X. Wang, B. Holcomb, R. H. Tien, D. K. Kim, L. Fenno, C. Ramakrishnan, et al. Cell-type-specific population dynamics of diverse reward computations. Cell 185, no. 19, 3568–3587.e27. (2022)

[46] C. Stringer, M. Michaelos, D. Tsyboulski, S. E. Lindo, and M. Pachitariu. High-precision coding in visual cortex. Cell 184, no. 10, 2767–2778.e15. (2021)

[47] W. Dabney, Z. Kurth-Nelson, N. Uchida, C. K. Starkweather, D. Hassabis, R. Munos, and M. Botvinick. A distributional code for value in dopamine-based reinforcement learning. Nature 577, no. 7792, 671–675. (2020)

[48] D. Gagnon and M. Parent. Distribution of VGLUT3 in highly collateralized axons from the rat dorsal raphe nucleus as revealed by single-neuron reconstructions. PLoS ONE 9, no. 2, e87709. (2014)

[49] Y. Dan, J. J. Atick, and R. C. Reid. Efficient coding of natural scenes in the lateral geniculate nucleus: Experimental test of a computational theory. The Journal of Neuroscience 16, no. 10, 3351–3362. (1996)

[50] N. Brenner, W. Bialek, and R. De Ruyter Van Steveninck. Adaptive rescaling maximizes information transmission. Neuron 26, no. 3, 695–702. (2000)

[51] K. Doya. Temporal difference learning in continuous time and space. Neural Information Processing Systems 8, 1073–1079. (1996)

[52] K. Beier, E. Steinberg, K. DeLoach, S. Xie, K. Miyamichi, L. Schwarz, X. Gao, E. Kremer, R. Malenka, et al. Circuit architecture of VTA dopamine neurons revealed by systematic input-output mapping. Cell 162, no. 3, 622–634. (2015)

[53] K. Miyazaki, K. Miyazaki, K. Tanaka, A. Yamanaka, A. Takahashi, S. Tabuchi, and K. Doya. Optogenetic activation of dorsal raphe serotonin neurons enhances patience for future rewards. Current Biology 24, no. 17, 2033–2040. (2014)

[54] M. Fonseca, M. Murakami, and Z. Mainen. Activation of dorsal raphe serotonergic neurons promotes waiting but is not reinforcing. Current Biology 25, no. 3, 306–315. (2015)

[55] J. N. McNair. Optimal giving-up times and the marginal value theorem. The American Naturalist 119, no. 4, 511–529. (1982)

[56] R. S. Sutton, D. Precup, and S. Singh. Between MDPs and semi-MDPs: A framework for temporal abstraction in reinforcement learn-ing. Artificial Intelligence 112, no. 1-2, 181–211. (1999)

[57] P. A. Correia, E. Lottem, D. Banerjee, A. S. Machado, M. R. Carey, and Z. F. Mainen. Transient inhibition and long-term facilitation of locomotion by phasic optogenetic activation of serotonin neurons. eLife 6, e20975. (2017)

[58] J. Qi, S. Zhang, H.-L. Wang, H. Wang, J. De Jesus Aceves Buendia, A. F. Hoffman, C. R. Lupica, R. P. Seal, and M. Morales. A glutamatergic reward input from the dorsal raphe to ventral tegmental area dopamine neurons. Nature Communications 5, no. 1, 5390. (2014)

[59] E. C. Azmitia. Chapter 1: Evolution of serotonin: sunlight to suicide in Handbook of Behavioral Neuroscience, pp. 3–22. C. P. Müller and K. A. Cunningham, eds. (Elsevier, 2020)

[60] J. C. Marques, M. Li, D. Schaak, D. N. Robson, and J. M. Li. Internal state dynamics shape brainwide activity and foraging behaviour. Nature 577, no. 7789, 239–243. (2020)

[61] T. E J. Behrens, M. W. Woolrich, M. E. Walton, and M. F S. Rushworth. Learning the value of information in an uncertain world. Nature Neuroscience 10, no. 9, 1214–1221. (2007)

[62] P. Piray and N. D. Daw. A model for learning based on the joint estimation of stochasticity and volatility. Nature Communications 12, no. 1, 6587. (2021)

[63] B. Weissbourd, J. Ren, K. E. DeLoach, C. J. Guenthner, K. Miyamichi, and L. Luo. Presynaptic partners of dorsal raphe serotonergic and GABAergic neurons. Neuron 83, no. 3, 645–662. (2014)

[64] B. A. Bari, C. D. Grossman, E. E. Lubin, A. E. Rajagopalan, J. I. Cressy, and J. Y. Cohen. Stable representations of decision variables for flexible behavior. Neuron 103, no. 5, 922–933.e7. (2019)

[65] M. Matsumoto and O. Hikosaka. Lateral habenula as a source of negative reward signals in dopamine neurons. Nature 447, no. 7148, 1111–1115. (2007)

[66] G. D. Stuber and R. A. Wise. Lateral hypothalamic circuits for feeding and reward. Nature Neuroscience 19, no. 2, 198–205. (2016)

[67] M. B. Lynn, S. Geddes, M. Chahrour, S. Maillé, E. Harkin, É. Harvey-Girard, S. Haj-Dahmane, R. Naud, and J.-C. Béïque. A slow 5-HT1AR-mediated recurrent inhibitory network in raphe computes contextual value through synaptic facilitation. bioRxiv. (2022) doi: 10.1101/2022.08.31.506056

[68] R. Wang and G. Aghajanian. Correlative firing patterns of serotonergic neurons in rat dorsal raphe nucleus. The Journal of Neuroscience 2, 11–16. (1982)

[69] S. D. Geddes, S. Assadzada, D. Lemelin, A. Sokolovski, R. Bergeron, S. Haj-Dahmane, and J.-C. Béïque. Target-specific modulation of the descending prefrontal cortex inputs to the dorsal raphe nucleus by cannabinoids. Proceedings of the National Academy of Sciences 113, no. 19, 5429–5434. (2016)

[70] D. Kim, H. H. Schütt, and W. J. Ma. Reward prediction error neurons implement an efficient code for reward. bioRxiv. (2022) doi: 10.1101/2022.11.03.515104

[71] M. Sousa, P. Bujalski, B. Cruz, K. Louie, D. McNamee, and J. Paton. Dopamine neurons reveal an efficient code for a multidimen-sional, distributional map of the future. Poster presented at COSYNE. (2023)

[72] E. F. Harkin, J.-C. Béïque, and R. Naud. A user’s guide to generalized integrate-and-fire models in Computational Modelling of the Brain: Modelling Approaches to Cells, Circuits and Networks, pp. 69– M. Giugliano, M. Negrello, and D. Linaro, eds. (Springer, 2021)

[73] E. F. Harkin, P. R. Shen, A. Goel, B. A. Richards, and R. Naud. Parallel and recurrent cascade models as a unifying force for understanding subcellular computation. Neuroscience 489, 200–215. (2022)

[74] C. Vandermaelen and G. Aghajanian. Electrophysiological and phar-macological characterization of serotonergic dorsal raphe neurons recorded extracellularly and intracellularly in rat brain slices. Brain Research 289, no. 1-2, 109–119. (1983)

[75] L. H. Calizo, A. Akanwa, X. Ma, Y.-z. Pan, J. C. Lemos, C. Craige, L. A. Heemstra, and S. G. Beck. Raphe serotonin neurons are not homogenous: Electrophysiological, morphological and neurochemical evidence. Neuropharmacology 61, no. 3, 524–543. (2011)

[76] K. Wong-Lin, G. Prasad, and T. M. McGinnity. A spiking neuronal network model of the dorsal raphe nucleus. The 2011 International Joint Conference on Neural Networks, 1591–1598. (2011)

[77] H. C. Tuckwell and N. J. Penington. Computational modeling of spike generation in serotonergic neurons of the dorsal raphe nucleus. Progress in Neurobiology 118, 59–101. (2014)

[78] K. Nakamura, M. Matsumoto, and O. Hikosaka. Reward-dependent modulation of neuronal activity in the primate dorsal raphe nucleus. The Journal of Neuroscience 28, no. 20, 5331–5343. (2008)

[79] E. S. Bromberg-Martin, O. Hikosaka, and K. Nakamura. Coding of task reward value in the dorsal raphe nucleus. The Journal of Neuroscience 30, no. 18, 6262–6272. (2010)

[80] Y.-Y. Feng, E. S. Bromberg-Martin, and I. E. Monosov. Dorsal raphe neurons signal integrated value during multi-attribute decision-making. bioRxiv. (2023) doi: 10.1101/2023.08.17.553745

[81] H. van Seijen and R. Sutton. True online TD(lambda). International Conference on Machine Learning 32, no. 1, 692–700. (2014)

[82] T. C. Hesterberg. What teachers should know about the bootstrap: Resampling in the undergraduate statistics curriculum. The American Statistician 69, no. 4, 371–386. (2015)

